# The developing tendon and enthesis are hypoxic and rely on hypoxia-inducible factor 1a (*Hif1a*) during postnatal development

**DOI:** 10.64898/2025.12.17.693209

**Authors:** Stephanie S. Steltzer, Nicole Migotsky, Tessa Phillips, Syeda N. Lamia, Ki Won Lee, Sueng-Ho Bae, Connor Leek, Sydney Grossman, Moaid Shaik, Allison Risha, Kaitlyn Frey, Claudia Loebel, Jun Hee Lee, Yatrik M. Shah, Adam C. Abraham, Megan L. Killian

**Author notes:** Correspondence: Megan L. Killian, PhD, North Campus Research Complex, 2800 Plymouth Road, Dock 90, Ann Arbor, MI 48109, United States.

## Abstract

The tendon-bone enthesis is a specialized fibrocartilaginous tissue crucial for muscle-to-bone force transmission, yet its postnatal development is not fully understood. Emerging evidence suggests hypoxia plays a pivotal role in enthesis maturation akin to its function in skeletal growth plates, with Hypoxia Inducible Factor 1 alpha (HIF-1α) acting as a key regulator of cellular adaptation (e.g., cell survival, extracellular matrix (ECM) deposition). Here, we investigated the spatial and temporal dynamics of hypoxia in the murine Achilles tendon enthesis and elucidated the role of Hif1a in enthesis cell survival and ECM formation using Scleraxis-lineage conditional knockout (ScxCre; Hif1a cKO) mice. We found that while neonatal tendons rapidly resolve hypoxia after birth, the enthesis maintains a hypoxic niche through postnatal day 5, mirroring a gradient observed in growth plates. Disruption of HIF-1α in enthesis-resident cells resulted in pronounced deficits in grip strength, abnormal tendon-bone attachment morphology, disrupted calcaneal architecture, impaired mineralization, and significant ECM disorganization. Histological analyses revealed persistent cell death and loss of the characteristic fibrocartilaginous gradient in cKO entheses, including dysregulated collagen alignment. In vitro, HIF-1α-deficient tendon fibroblasts exhibited blunted transcriptional responses to hypoxia, altered metabolic gene expression, and changes in ECM deposition. Collectively, our findings illuminate hypoxia as a sustained niche in the postnatal enthesis, with HIF-1α critically required for cell survival, ECM organization, and enthesis structural integrity. This work advances our understanding of enthesis biology and provides insights relevant to tendon-bone attachment disorders and regenerative strategies.

## Introduction

The tendon-bone attachment (i.e., enthesis) is essential for the transmission of muscle forces to bone.^1,2^ The progenitor pool that forms the enthesis also contributes to tendon development and is identified by the transcription factor, *Scleraxis* (*Scx*).^3–5^ The enthesis develops analogously to the skeletal growth plate of long bones where cartilage is replaced with bone via endochondral ossification.^6–8^ However, while the growth plates eventually fuse, the fibrocartilage enthesis remains static throughout the lifespan and has been likened to an “arrested” growth plate.^9^

During embryonic development, the growth plates of long bones maintain a hypoxic niche that resolves with vascularization.^10–12^ *Hypoxia inducible factor 1 alpha* (HIF-1α) is a master transcriptional regulator of cell survival in hypoxic environments and modulates metabolic pathways that promote the deposition of extracellular matrix (ECM) and vascularization.^13,14^ HIF-1α is required for the survival of growth plate chondrocytes during development, and loss of *Hif1a* results in cell death and structural abnormalities in developing bone.^11,14^ However, the enthesis is well known to maintain limited vascularization, suggesting that cellular hypoxia may play a sustained role in its maintenance.

Hypoxia also regulates cell proliferation, and tendon fibroblasts are highly proliferative during embryonic tendon development prior to when high amounts of ECM are deposited. However, the cells in the mature tendon do not retain the same level of proliferation.^15^ Likewise, the density of resident cells in the postnatal enthesis decreases as its resident cells secrete and remodel their ECM, with little to no proliferation during this time.^16^ The enthesis consists of both cellular and ECM gradients from tendon to bone that are established during post-natal growth.^4,9,17^ The gradient-like distribution of cells in the enthesis as well as tendon may be driven by changes in the oxygen environment, yet this spatial distribution of hypoxia has been unclear until now.

In this study, we hypothesized that hypoxia is sustained in postnatal enthesis and that enthesis-resident cells rely on HIF-1α for their survival and ECM deposition. We first explored if and when enthesis-resident cells experience hypoxic stress during embryonic and neonatal growth in mice. We then identified a critical role of *Hif1a* for enthesis cell survival, ECM organization, spatial transcriptome, and biomechanical function in the mouse Achilles tendon and enthesis with targeted deletion of HIF-1α in the *Scx*-lineage *in vivo*. Because the enthesis is composed primarily of Scx-lineage fibroblasts, we further investigated if and how murine fibroblasts respond *in vitro* to low oxygen environments. To do this, we measured the transcriptional response as well as cell and ECM dynamics of fibroblasts in hypoxia or following loss of *Hif1a in vitro*.

## Results

### Enthesis, but not tendon, remains hypoxic during early postnatal development

Due to the intrinsic link between hypoxia and HIF-1α, we first determined if a hypoxic environment exists at the embryonic and neonatal Achilles tendon-bone enthesis using EF5 staining, which forms intracellular adducts in hypoxic conditions (<10% O_2_) that can be visualized via immunostaining. EF5 staining for hypoxic cells revealed that both embryonic tendon and enthesis experience hypoxia, with strong EF5 staining at E16.5 and P1 (Fig. 1). Additionally, the enthesis remained hypoxic until P5 while the developing tendon was no longer experiencing hypoxic stress at P5 (Fig. 1).

**Figure 1.**
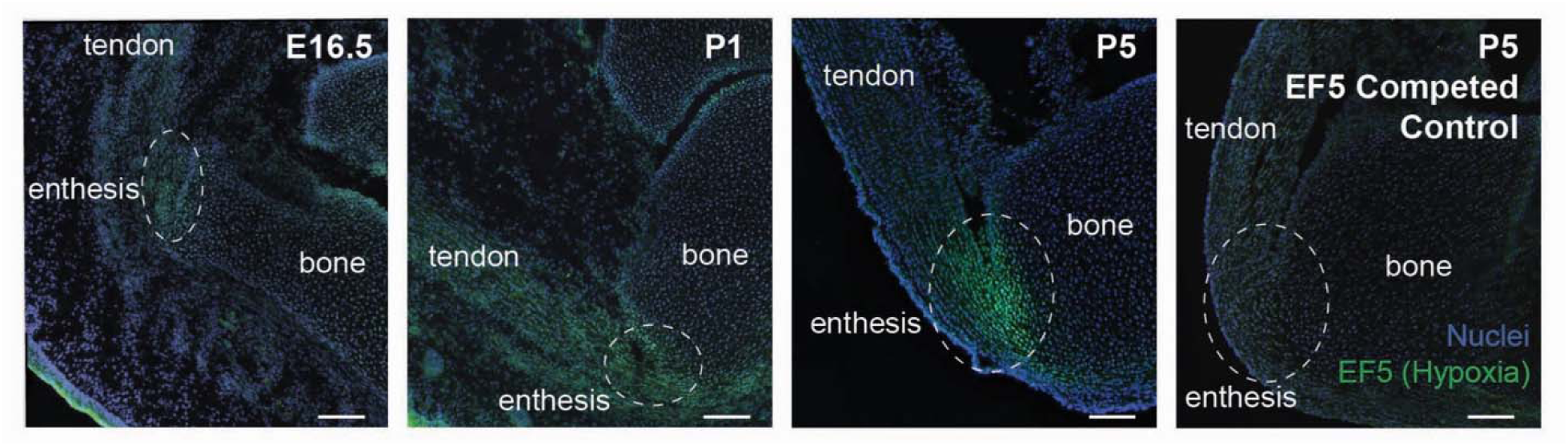
Enthesis, but not tendon, remains hypoxic during post-natal growth. Accumulation of EF5 (hypoxia marker, green fluorescence) becomes more localized to the C57BL6/J mouse Achilles enthesis during early postnatal development (Embryonic day 16.5 (E16.5) through postnatal days 1 and 5 (P1 and P5)) compared to tendon and bone. EF5, green fluorescence; Nuclei, blue fluorescence. Scalebar = 200μm. Last panel is non-competed control.

### Loss of *Hif1a* in tendon progenitors leads to impaired grip force and deficient tendon/enthesis function

We investigated the Achilles tendon and enthesis of mice with or without *Hif1a* at several timepoints throughout development (Fig. 2A). In mice lacking *Hif1a* in *Scleraxis*-Cre (*Scx*Cre) cells (*Hif1a ^ScxCre^*; cKO), overall tendon patterning appeared grossly normal; however, tendon fusion was disrupted compared to 8-week (postnatal day 56, P56) littermate controls (Ctrl) which was visualized using optical coherence tomography with polarization (OCT) (Fig. 2B and Fig. 3). Additionally, forelimb grip strength was significantly lower for cKO mice compared to Ctrl, regardless of sex (Fig. 2C), while Ctrl and cKO mice were comparable in weight (Supplemental Fig. S1). The skeletal morphology, visualized using planar X-ray, showed that joint congruency in cKO mice were severely impaired, especially at sites where tendons attach to bone (e.g., the calcaneus of the heel and olecranon of the elbow, Fig. 2D). Additionally, the Achilles tendon length, width, and volume were comparable between adult Ctrl and cKO mice. However, we visualized using OCT that cKO Achilles tendons failed to fuse and remained bifurcated distally compared to Ctrl tendons by 8wks of age (Fig. 3). We evaluated calcaneal morphometry using nanoCT and found that cKO calcanei had incomplete and irregular bone formation adjacent to the Achilles enthesis in both male and female mice (Fig. 4A). While Ctrl mice did not present with any mineralization in their tendons by 8wks of age, we observed mineral accumulation in male, but not female, cKO mice at the distal Achilles tendon (Fig. 4A; 0.024 ± 0.013 mm^3^). Calcaneal bone volume ratio (BV/TV) was also reduced in cKO mice compared to Ctrl (Fig. 4B). Additionally, cKO calcanei exhibited notable morphological disruptions and were both shorter and wider than Ctrl calcanei (Fig. 4C and D). The widening of the calcanei in cKO mice was especially pronounced near the calcaneal tubercle (Supplemental Fig. S4) and accounted for 51.1% of the shape variation compared to Ctrl calcanei. Surprisingly, these major differences in bone and tendon structure did not translate to significant differences in biomechanical strength of the Achilles tendons, as adult Ctrl and cKO tendons did not show differences in maximum (failure) force or stress (Fig. 2E and F). However, linear stiffness (but not linear modulus) was significantly lower in cKO compared to Ctrl tendons, and the cross-sectional area of the cKO tendons was also smaller (p=0.0638) compared to Ctrl tendons (Fig. 2F), which did not translate to differences between groups in CSA-normalized stiffness (i.e., linear modulus; Fig. 2F). Despite this, the failure modes of tendons during biomechanical testing were different between Ctrl and cKO groups, in which all Ctrl Achilles tendons failed at the growth plate and all cKO tendons failed at either the Achilles enthesis or tendon midsubstance (Supplemental Fig. S2). This difference in failure modes was possibly due to poor bone quality at the enthesis in cKO mice and preservation of the calcaneal growth plate in the Ctrl but not cKO heels. The significant differences in tendon bifurcation, visualized using OCT, led us to further explore the role of HIF1a in the tendon and enthesis organization during key developmental stages, when new ECM is being deposited and remodeled.^17,18^

**Figure 2.**
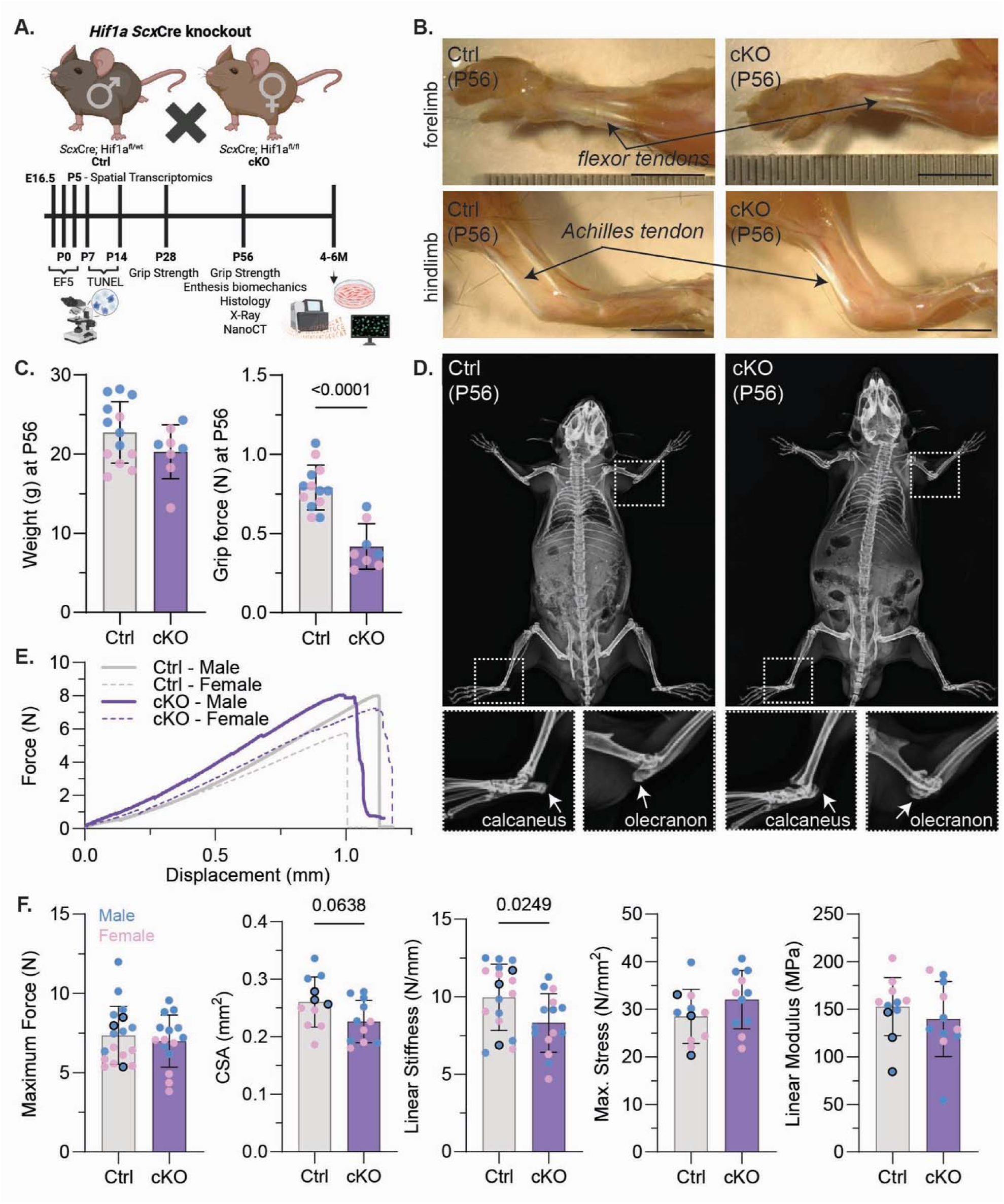
Loss of *Hif1a* in tendon leads to impaired grip force and deficient tendon function. (A) The murine knockout model of *Hif1a* in Scleraxis (SCX) lineage cells (*Hif1a^flox/flox^*;*ScxCre*; cKO) inhibits the production of *Hif1a*. (B) Force-generating tendons pattern normally in mice lacking *Hif1a* in tendons. Morphology of the forelimb (e.g., flexor tendons) and hindlimb (e.g., Achilles tendon) tendons in cKO mice were unremarkable compared to control (Ctrl) mice at postnatal day (P)56. Scalebar = 0.5mm. (C) Knockdown of *Hif1a* did not affect mouse weight at P56. Both male (blue) and female (pink) cKO mice had reduced forepaw grip strength. Two-way ANOVA (sex/genotype) with Dunnett corrected multiple comparisons. (D) Planar X-rays of cKO mice at P56 show marked disruptions in tendon-bone attachment sites compared to control, most notably in the elbow (olecranon, white dashed line) and ankle (calcaneus; white dashed line). (E) Representative force vs displacement curve for P56 Ctrl and cKO Achilles tendon-enthesis. (F) Ultimate load (N) of Achilles tendon-entheses at P56 was similar in cKO mice compared to control. Cross sectional area (CSA, mm) of Achilles tendons was reduced in cKO mice compared to control. Achilles tendon-enthesis stiffness (N/mm) was decreased in both male and female cKO mice compared to control, while maximum stress and linear modulus remained similar across genotypes. One-way ANOVA with Dunnett’s corrected multiple comparisons. Points outlined in black indicate *Scx*Cre-positive control animals.

**Figure 3.**
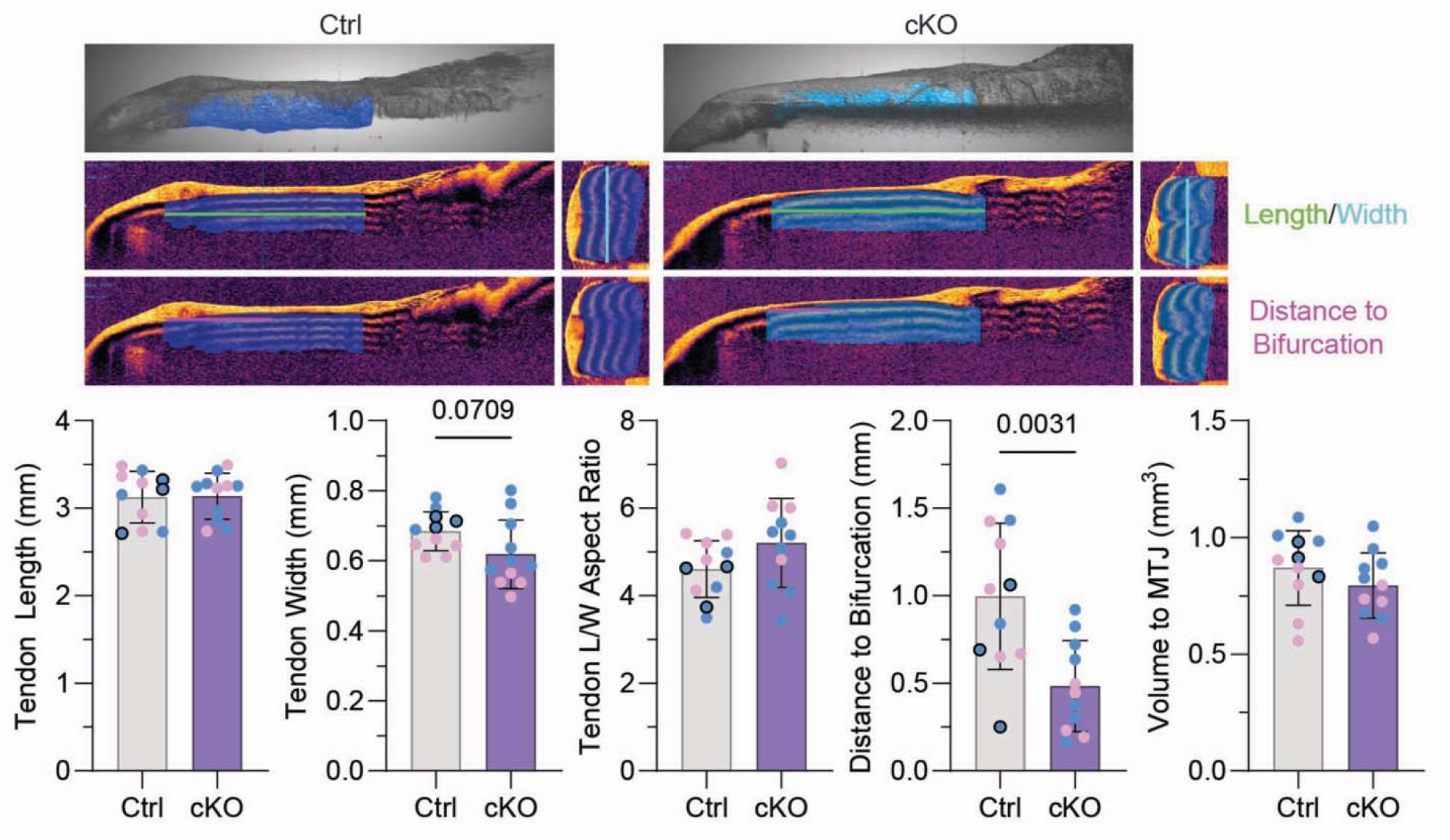
No changes were exhibited in Achilles tendon length and width while the distance to the tendon bifurcation were decreased in cKO mice compared to Ctrl. Representative images of Achilles tendon masks generated in Dragonfly ORS using retardation and intensity images. Achilles tendon length, width, and distance to bifurcation were quantified using optical coherence tomography (OCT). Blue indicates mask of tendon.

**Figure 4.**
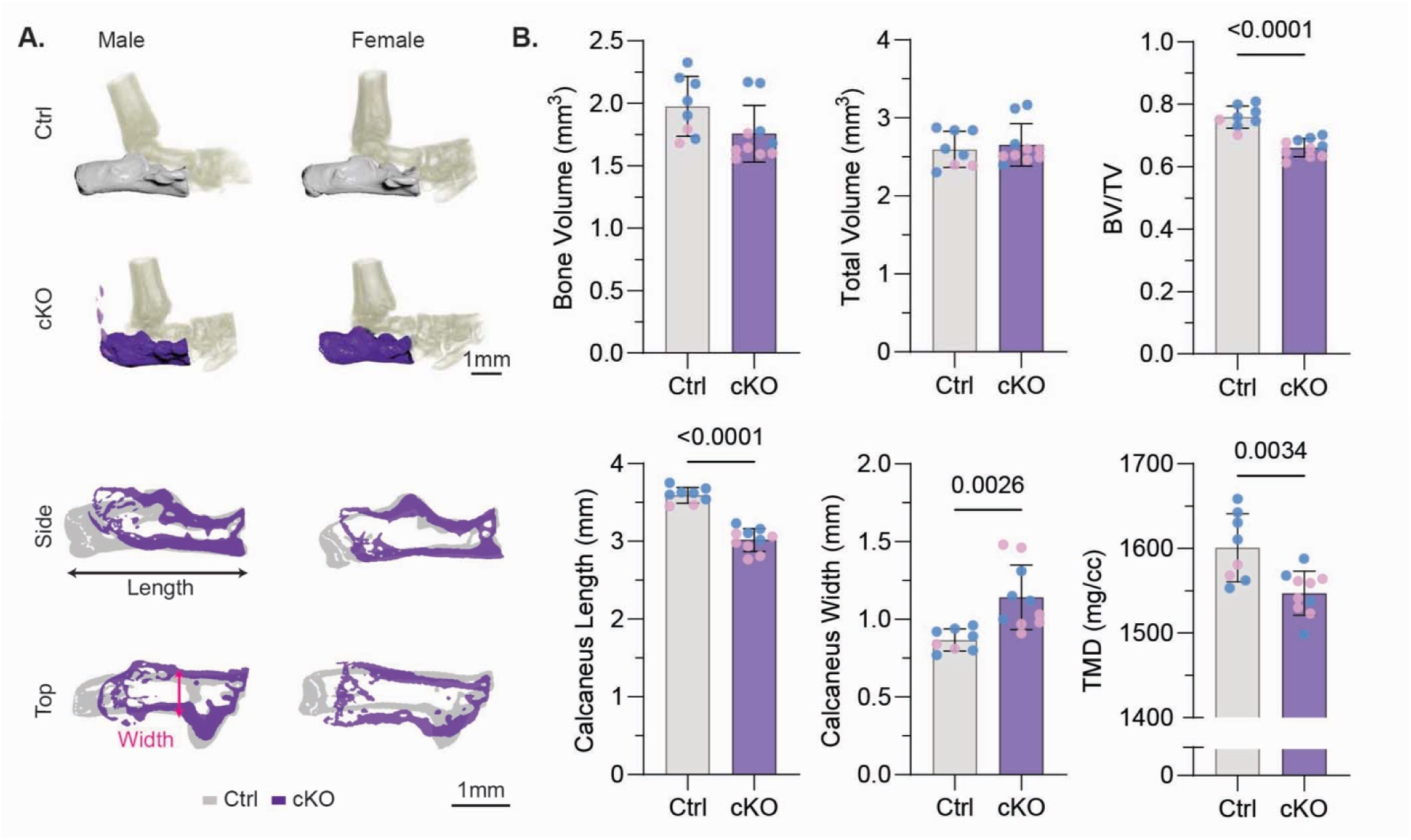
Loss of *Hif1a* disrupts calcanei shape and size at the Achilles tendon insertion site. (A) 3D and 2D nanoCT reconstructions of both male and female postnatal day (P)56 cKO calcanei revealed HIF-1α is required to maintain proper calcaneus shape and size near the Achilles tendon insertion site when compared to the control. Scalebar = 1mm. (B) NanoCT analyses of the calcaneus revealed a decrease in bone volume fraction (BV/TV) and calcaneus length of P56 cKO mice compared to control with an increase in calcaneus width in cKO mice compared to control. Tissue mineral density (TMD) also decreased in cKO mice compared to control. Mineralization was evident in the male cKO Achilles tendons. One-way ANOVA with Dunnett’s corrected multiple comparisons.

### Loss of *Hif1a* disrupts ECM organization and cell viability in the postnatal enthesis

Normal fibrocartilage development occurs postnatally in the mouse enthesis, with a postnatal mineralization and transition between mineralized and unmineralized fibrocartilage established around 2 weeks of age (∼P14).^9^ In the normally developing Achilles enthesis, we observed the cellular gradient by P7 and the transitional ECM between tendon and cartilage template by P14 (Fig. 5A). The fully mature Achilles enthesis was established by P56 (Fig. 5A and Supplemental Fig. 5A). In the cKO enthesis, the tendon-bone enthesis is established at P7, however by P14 and through P56, the Achilles tendon and enthesis began to exhibit more cell and ECM disorganization (Fig. 5A and Supplemental Fig. S3A). By P56, the deformity at the enthesis was characterized by focal bone loss proximal to the attachment site, focal regions of hypo- and hypercellularity in the insertional tendon, and ECM disorganization at the tendon-bone interface (Fig. 5A and Supplemental Fig. 3A). We also found that cell death using TUNEL staining was consistent between genotypes at P7, cell death remained elevated in cKO but not Ctrl entheses by P14 (Fig. 5B) and resolved to comparable levels by P56 (Supplemental Fig. 3B). This suggests that *Hif1a* plays a critical role in enthesis cell survival during postnatal maturation of the fibrocartilage enthesis.

**Figure 5.**
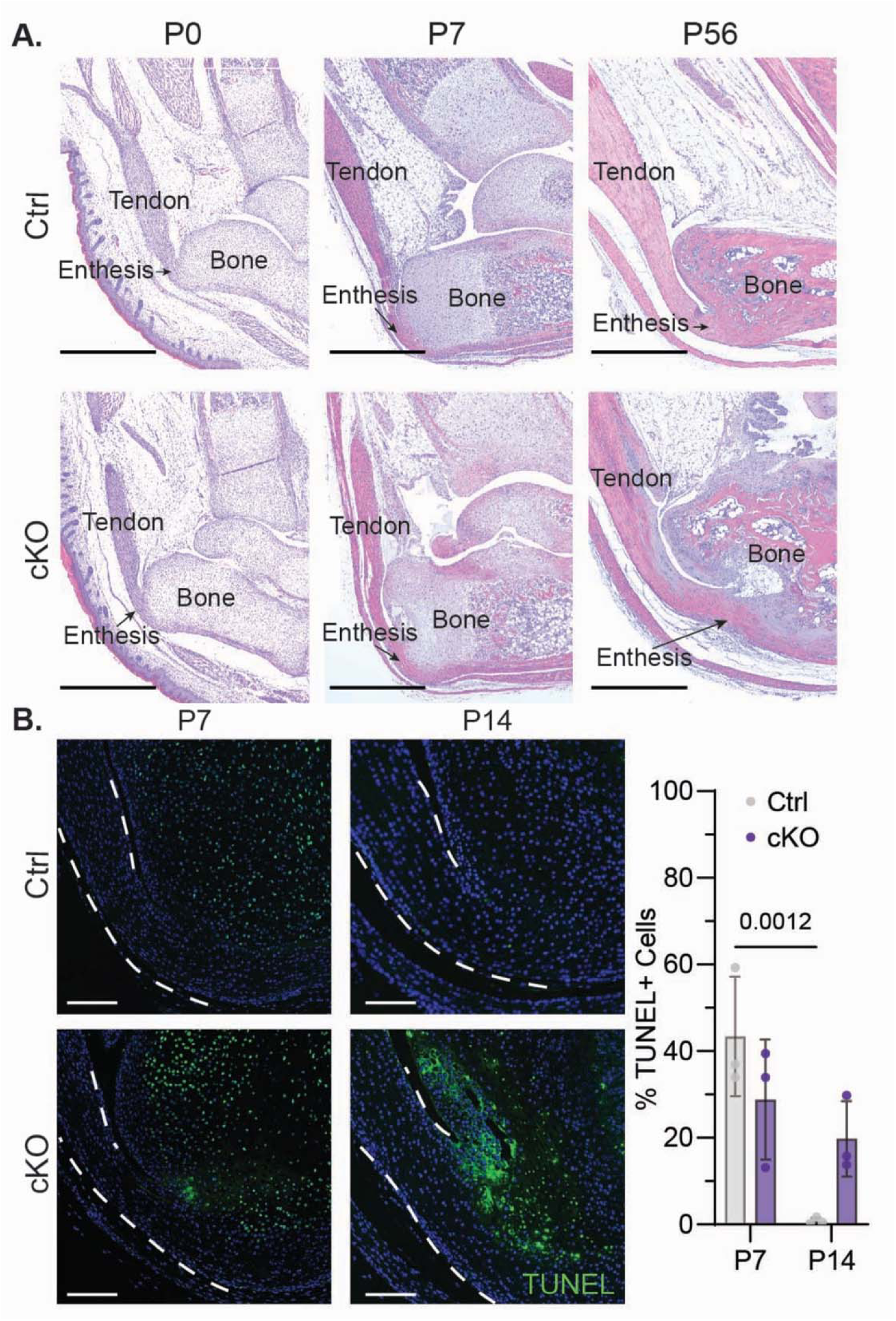
The developing tendon and enthesis are reliant on *Hif1a* for structure and cell survival. (A) Hematoxylin and Eosin staining at P0 show unremarkable differences due to *Hif1a* knockout while disorganized cell and ECM are present at P7 and P56 entheses. Scalebar = 1mm. (B) Cells in the insertional tendon (white dashed line) and enthesis exhibit more DNA damage and cell death (TUNEL+ cells, green fluorescence) in P14 cKO mice when compared to control. Two-way ANOVA with Tukey’s multiple comparisons test.

### *Hif1a* is required for the formation of a structural gradient at the Achilles enthesis

Using Silver nitrate staining, which is commonly used to stain osteocytes in bone,^19^ we observed a well-defined tidemark that separates mineralized and unmineralized fibrocartilage in the Ctrl Achilles enthesis by P56 (Fig. 6A). However, in age-mateched mice, cKO entheses had diffuse Silver+ staining into the insertional tendons that was not present in Ctrl tendons (Fig. 6A and B). Additionally, the distinct tidemark was not present in cKO entheses (Fig. 6A). We also observed, using Safranin O staining for acidic proteoglycans and glycosaminoglycans (commonly used to stain cartilage), that Ctrl insertional tendons were sparse in proteoglycan-rich cells while cKO tendons had significantly more proteoglycan-rich cells adjacent to the articular surface (Fig. 6B). Additionally, cKO insertional tendons had more cells (higher cell density) compared to Ctrl tendons at P56 (Fig. 6B). This may be caused by a localized shift in tendon compression in cKO mice.^8,20,21^ We also saw a severe disruption and loss of Safranin-O+ growth plates of cKO mice, which remained distinct in Ctrl calcanei at P56 (Fig. 6B). To further interrogate ECM organization, we used Picrosirius Red staining and quantitative polarized light imaging (qPLI) to quantify collagen alignment in the insertional tendon and found Ctrl tendons had highly aligned collagen at P56 (Fig. 6C). However, cKO tendons were less uniformly aligned, as quantified as reduced average degree of linear polarization (DoLP). Taken together, these data suggest that survival of cells within and adjacent to the enthesis is necessary to generate the gradient ECM needed for proper enthesis maintenance.

**Figure 6.**
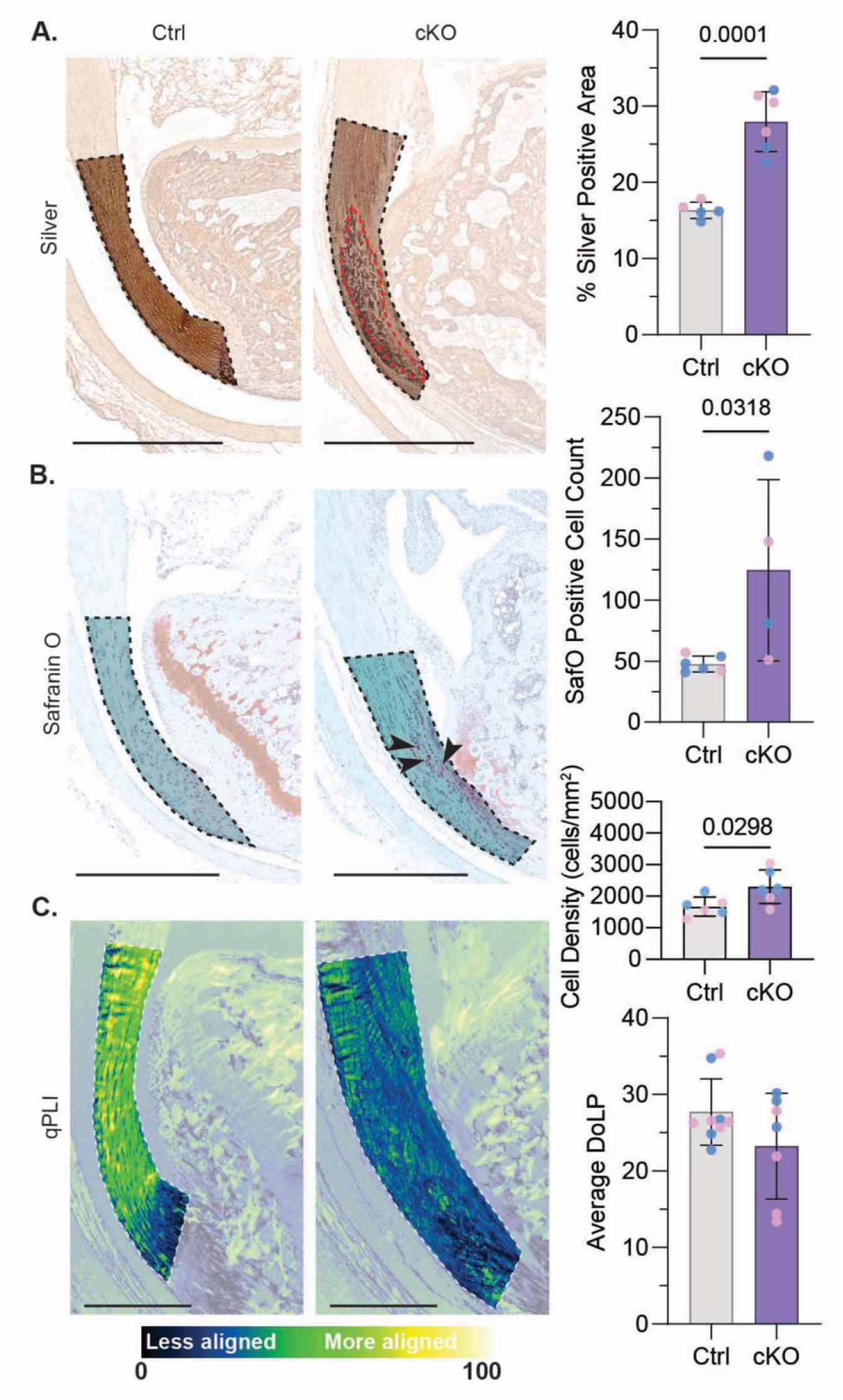
*Hif1a* is required for structural integration of the tendon into the bone. (A) Silver Nitrate sections of cKO mouse Achilles enthesis contained a build-up of DNA and protein compared to the age-matched Cre-control (Ctrl) at P56. Increased areas of protein accumulation (red dashed line) occurred in cKO mouse entheses with poorly defined secondary ossification center compared to Ctrl. (B) Safranin O sections of P56 cKO mouse Achilles enthesis revealed cKO growth plates were poorly organized and an increased number of proteoglycan-rich cells (black arrows) were apparent in the cKO insertional tendon when compared to Ctrl. Total cell density within the insertional tendons was significantly higher for cKO tendons compared to Ctrl. (C) Degree of linear polarization images of Achilles insertional tendon identified decreased collagen organization in P56 cKO mice when compared to Ctrl. Scalebar = 1mm. One-way ANOVA with Dunnett’s corrected multiple comparisons.

### Knockdown of *Hif1a* in mouse tail tendon fibroblasts (TFs) desensitizes TFs to hypoxic conditions (1% O_2_) compared to normoxic conditions (20% O_2_)

Due to the established hypoxic gradient at the enthesis and the clear disruption in tendon and enthesis organization in cKO mice, we then investigated the role of HIF-1α in regulating oxygen-dependent transcriptional programs and ECM in TFs. We found that mouse tail TFs from *Hif1a* cKO mice had reduced sensitivity to hypoxia after 1-week in culture, as indicated by dampened expression of hypoxia-sensitive genes (e.g., *Hilpda*; Fig. 7A). In both Ctrl and cKO TFs, we found that 1% O_2_ led to downregulation of mechanoresponsive genes, *Scx* and *Igfbp9* (Fig. 7A). To define the extent to which hypoxia and *Hif1a* loss reshape TF transcriptional programs, we performed bulk RNA-sequencing after 4 days in culture. In 1% O_2_, Ctrl TFs were enriched for genes associated with metabolic switching (Fig. 7B and C), including the HIF-1 signaling pathway (24 out of 708 differentially expressed genes; log_2_ fold change of 2.77) and glycolysis and gluconeogenesis (19 genes; log_2_ fold change of 3.83), and the pentose phosphate pathway (11 genes; log_2_ fold change of 4.37). We also identified enrichment of molecular pathways associated with ECM deposition and remodeling, including osteoclast differentiation (*Osr2, Runx2, Smad6, Smad7, Smad9, Bmp4, Tgfbr1, Tgfbr2*), proteoglycans, and regulation of the actin cytoskeleton (Fig. 7B and C, and Supplemental Fig. S5).

**Figure 7.**
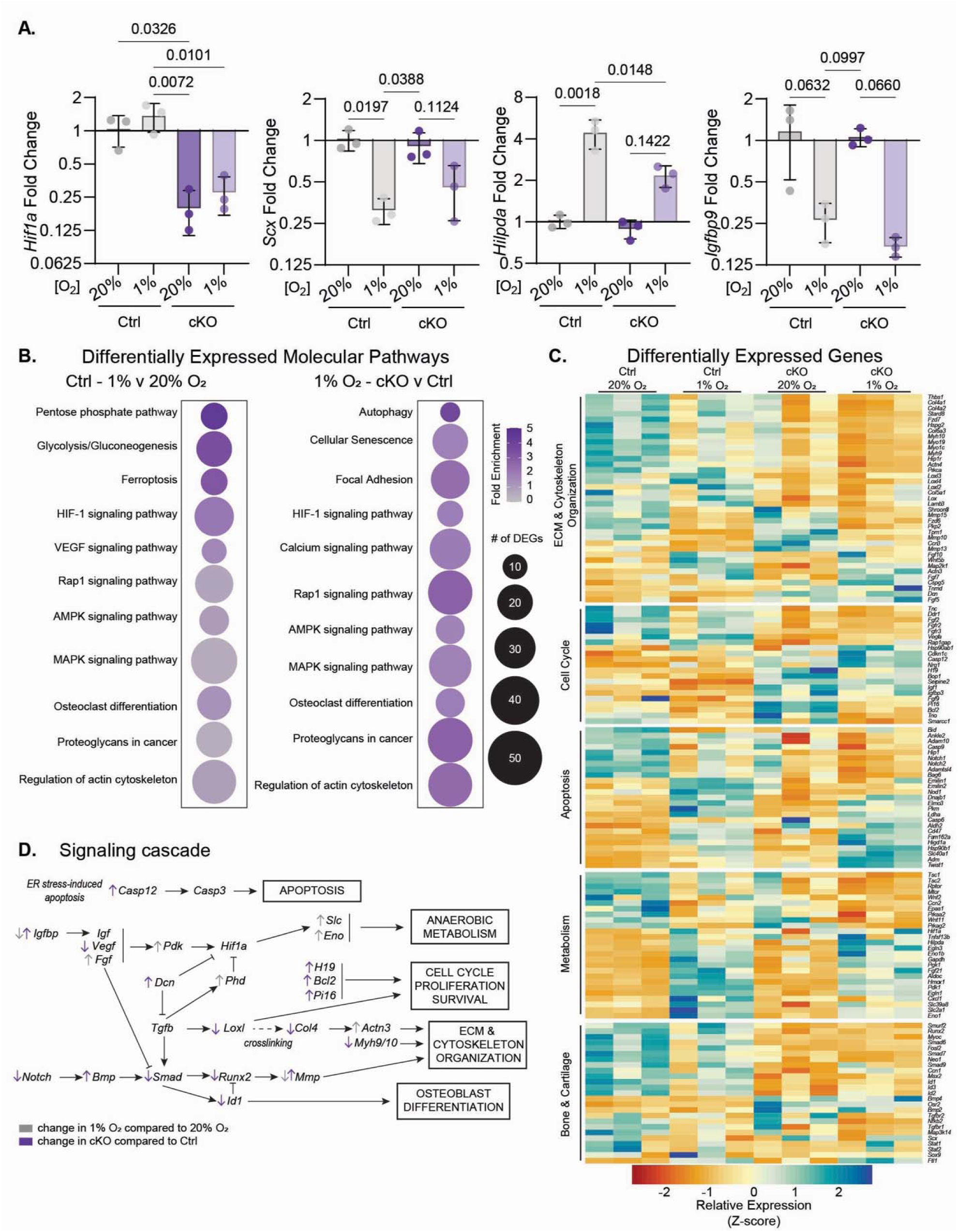
Knockdown of *Hif1a* in mouse tail tendon fibroblasts (TFs) desensitizes TFs to hypoxic conditions (1% O_2_) compared to normoxic conditions (20% O_2_). (A) *Hif1a* relative expression is reduced in mouse tail tendon fibroblasts cultured from mice lacking *Hif1a*. TFs cultured for 1-week in hypoxia express reduced tenogenic (*Scx*) and mechanotransductive (*Igfbp9*) genes as well as activated hypoxia-related genes (*Hilpda*). Each dot represents a biological replicate; (2^-ΔCq^; *Actb*, *Polr2a* reference genes). Bars show mean ±SD of ΔC_q_; Two-way ANOVA tests with Dunnett’s multiple comparisons for hypoxia and genotype. (B) Quantification of Kegg pathways found during bulk RNA sequencing in Ctrl 1% O_2_ compared to Ctrl 20% O_2_ and cKO 1% O2 compared to Ctrl 1% O_2_. (C) Heatmap depicting DEGs affiliated with ECM and cytoskeleton organization, cell cycle, apoptosis, metabolism, and bone and cartilage. Differentially expressed genes were determined using a fold change cutoff of 1.5 and a p-value cutoff of 0.05. (D) Signaling diagram created from DEGs.

In cKO TFs, *Casp12* was elevated compared to Ctrl regardless of oxygen availability, suggesting endoplasmic reticulum stress-induced apoptosis as a potential mechanism of cell death in the absence of *Hif1a* (Fig. 7D). In 1% O_2_, there were no differentially expressed caspase genes compared to 20% O_2_. Additionally, cKO TFs had upregulated expression of genes associated with the cell cycle, proliferation, and survival including the long non-coding RNA, *H19,* and the apoptosis inhibitor, *Bcl2*, as well as *Pi16*, a marker of fibroblast specification (Fig. 7D). Even in cKO cells, TFs cultured in 1% O_2_ had elevated genes associated with metabolism (e.g., *Slc40a1* and *Eno1*) compared to 20% O_2_, suggesting that *Hif1a* is not required for glycolytic and iron metabolism in hypoxia *in vitro* (Fig. 7D).

We found that, in cKO TFs, both *Loxl* and *Col4* expression were suppressed, as was *Myh9/10*, suggesting a potential role for *Hif1a* in ECM crosslinking and cytoskeletal organization (Fig. 7D). *Bmp4* was upregulated in cKO TFs compared to Ctrl TFs, while *Notch*, *Smad*, *Runx2*, and *Id1* were suppressed (Fig. 7D). This suggests that cKO cells are prevented from fully differentiating into bone-forming osteoblasts.^22,23^ Despite this, cKO cells still generate a matrix consistent with disorganized pathogenic mineralization, indicating a process of calcification or heterotopic ossification.^22,23^

### Hypoxic (1% O_2_) conditions *in vitro* leads to dysregulation in fibroblast metabolism and nascent matrix deposition regardless of genotype

After 4 days in culture, we observed decreased ATP availability in hypoxic conditions regardless of *Hif1a* expression (when normalized to cell density), supporting that TFs in hypoxia do not produce as much ATP as TFs in normoxia and instead rely on glycolysis (Fig. 8). Cell density did not significantly change at either the 2-day or 4-day time points, while ATP and matrix data at 2-days followed similar trends to 4-day cultures (Supplemental Fig. S6A and C). Nuclear circularity decreased in Ctrl cells cultured in hypoxia compared to normoxia while cKO cells did not exhibit a change in nuclear circularity, suggesting *Hif1a* is, in part, required for TF nuclear response to hypoxia (Fig. 8). Newly synthesized and secreted (nascent) matrix area was increased, although not significantly, in cKO cells cultured in normoxia compared to Ctrl cells, suggesting the cKO cells may generate more ECM under normoxic compared to hypoxic conditions (Fig. 8). Lastly, hypoxic conditions increased calcium and reduced GAG in addition to reducing ATP availability at 2-weeks (Supplemental Fig. S6B). To determine what transcriptional differences may lead to the structural and mechanical defects seen in the cKO tendon-to-bone interface, we performed spatial transcriptomics on postnatal day 5 (P5) Achilles entheses.

**Figure 8.**
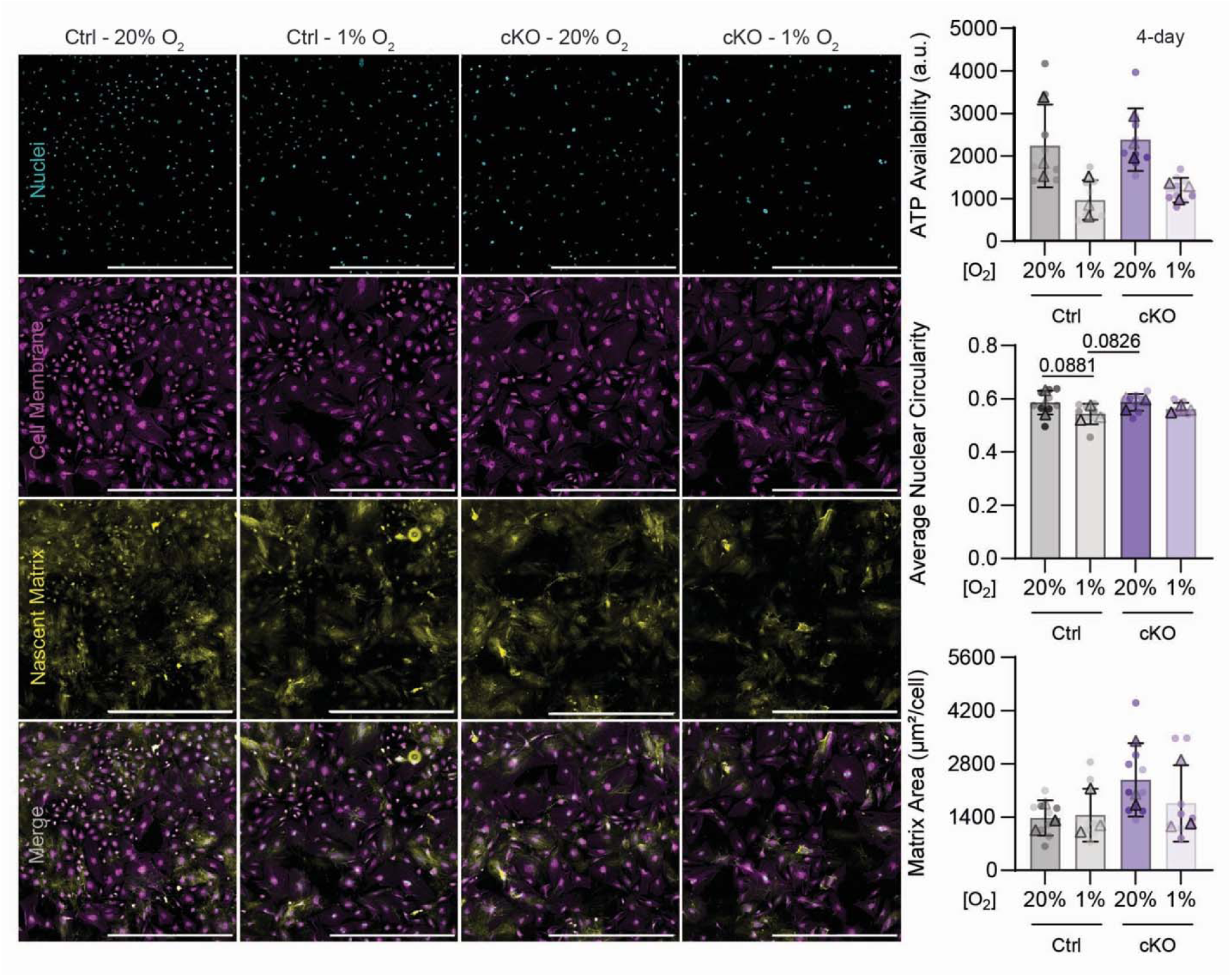
Knockdown of *Hif1a in vitro* causes dysregulation in metabolism and nascent matrix deposition. Nascent matrix staining of Ctrl and cKO TFs cultured in 20% O_2_ or 1% O_2_ for 4 days. Decreased ATP was available in hypoxic conditions regardless of genotype when normalized to cell density. Nuclear circularity decreased in Ctrl cells cultured in hypoxia compared to normoxia while cKO cells did not exhibit a change in nuclear circularity. Nascent matrix area may increase in cKO cells cultured in normoxia compared to other conditions when normalized to cell density. Bars show mean ±SD; Two-way ANOVA tests with Dunnett’s multiple comparisons for hypoxia and genotype. Triangles denote average of one biological replicate across 3 technical replicates.

**Table 1.** Readouts from *in vitro* data (mean ± standard deviation (S.D.)). *4-day* data: Cell Density, Total ATP, Total Nascent Matrix. *2-week* data: Extracellular calcium and Total GAG.

|  |  | 4-day |  |  | 2-week |  |
| --- | --- | --- | --- | --- | --- | --- |
|  | Oxygen | Cell density (number/mm<sup>2</sup>) | Total ATP availability (a.u.) | Total nascent matrix area (μm<sup>2</sup>/cell) | Extracellular calcium (μg/dl) | Total GAG from cell lysate (ng) |
| Control | 20% O<sub>2</sub> | 63 ± 27 | 2236 ± 812 | 1376 ± 289.8 | 5.39 ± 0.34 | 27.61 ± 0.02 |
|  | 1% O<sub>2</sub> | 63 ± 25 | 968 ± 397 | 1450 ± 499 | 7.59 ± 1.59 | 27.52 ± 0.01 |
| *Hif1a*<br>cKO | 20% O<sub>2</sub> | 44 ± 9 | 2384 ± 405 | 2380 ± 735 | 4.78 ± 0.27 | 27.59 ± 0.06 |
|  | 1% O<sub>2</sub> | 52 ± 10 | 1199 ± 168 | 1761 ± 805 | 7.47 ± 1.87 | 27.52 ± 0.01 |

### Spatial sequencing at postnatal day 5 (P5) showed probable cell cycle dysregulation in cKO Achilles tendon and bone, with cKO tendon more similar to cKO bone than control tendon is to control bone

Postnatal day 0 (P0) cKO tendon-to-bone interfaces appeared to pattern similarly to Ctrl based on histology with defects showing by P7 (Fig. 5A, Supplemental Fig. S3). Therefore, sections from P5 Achilles and entheses were used to compare transcriptional differences between Ctrl and cKO mice that may lead to the dysregulation of cell and ECM organization. The spatial transcripts were first segmented into insertional tendon and bone regions (Fig. 9A). We found a clear divergence in spatial patterning for Ctrl entheses but not for cKO entheses, with more differentially expressed genes and pathways identified for comparisons of Ctrl tendon to bone versus cKO tendon to bone (Fig. 9B-E). That more genes diverge spatially in Ctrl entheses suggests that the insertional tendon of the cKO Achilles may share similarities with the cKO bone, whereas the Ctrl tendon and bone have, at this time point, already diverged transcriptionally from each other (Fig. 9B). Further, within the pathways unique to the Ctrl tendon vs. bone comparison was the response to hypoxia, and inhibition of non-skeletal tissue mineralization (Fig. 9B). Pathways unique to the cKO tendon vs. bone comparison indicated changes in the regulation of cell cycle and cell growth, primarily in response to DNA damage during the G1 to S transition (Fig. 9B). Although there were some similarities in the differentially expressed genes in tendon and in bone regardless of genotype (e.g., *Col1a1, Col1a2, Fmod* upregulated in tendon for both Ctrl and cKO, and *Col2a1*, *Col3a1*, *Col9a1-3*, *Col11a2*, *H19* upregulated in bone for both Ctrl and cKO), there were a reduced number of genes associated with these pathways for the cKO sample compared to Ctrl (Fig. 9C and D). Fewer DEGs were identified in the oxidative phosphorylation, AMPK signaling, and proteoglycans in cancer pathways. Finally, we found that several of the genes downregulated in cKO tendon compared to Ctrl tendon were associated with skeletal system morphogenesis, proliferation, and regulation of cell cycle and cell growth while genes associated with iron homeostasis, filament organization, and response to oxidative stress were upregulated in cKO tendon compared to Ctrl (Fig. 9E). These data indicate disruption of crucial cell survival and tissue delineation processes in the cKO tendons compared to Ctrl.

**Figure 9.**
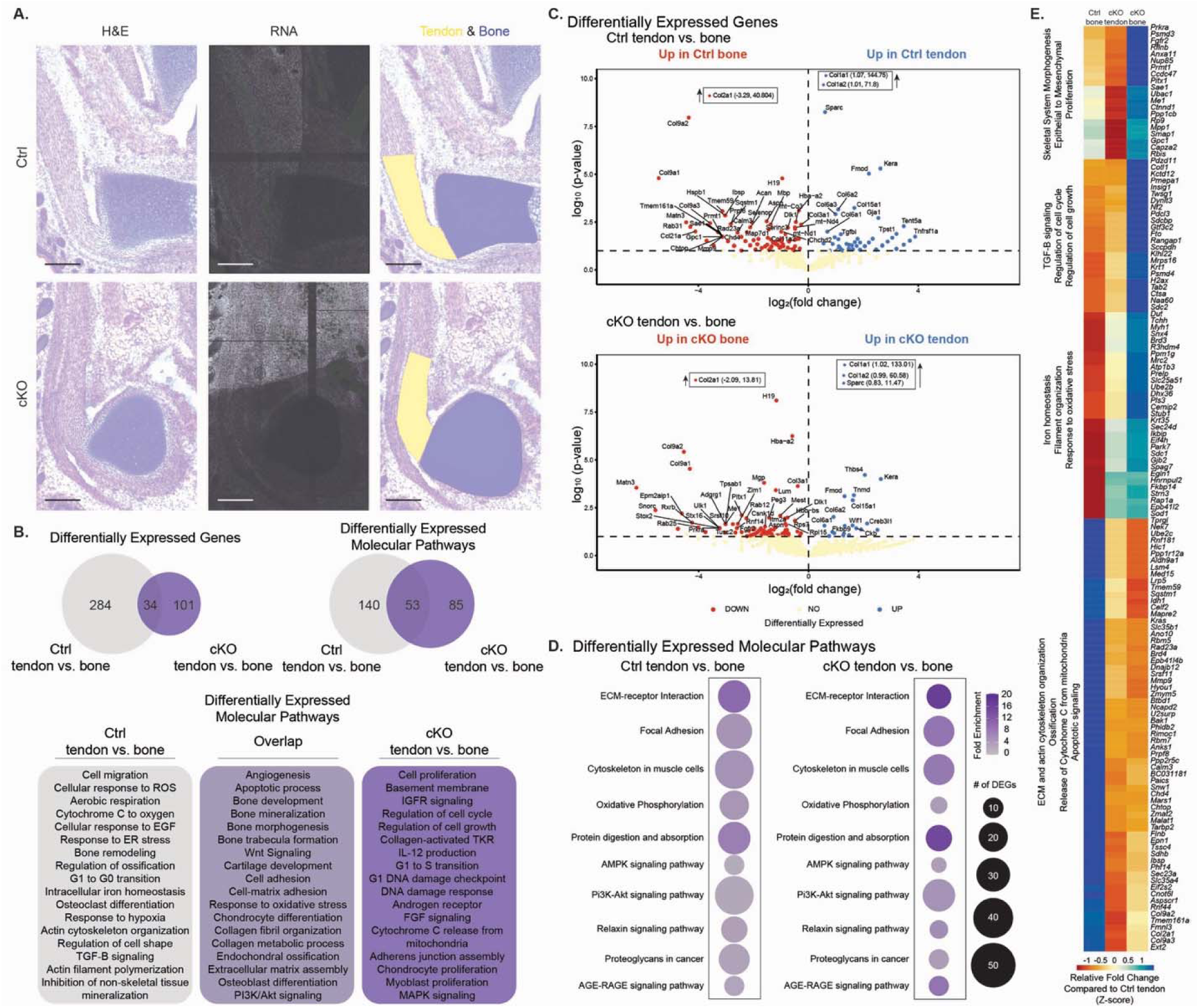
Spatial sequencing at postnatal day 5 (P5) showed probable cell cycle dysregulation in cKO Achilles tendon and bone, with cKO tendon more similar to cKO bone than control tendon is to control bone. (A) Hematoxylin and Esoin stained sections of P5 mouse Achilles tendon insertions with transcripts overlapped and the tendon (yellow) and bone (blue) segmented; Scalebar = 200μm. (B) Differentially expressed genes (DEGs) and molecular pathways in Ctrl tendon vs. bone comparison and cKO tendon vs. bone comparison. (C) Volcano plots of DEGs as defined by a p-value cutoff of 0.1. Top 50 DEGs labeled. (D) Differentially expressed molecular pathways in Ctrl tendon vs. bone and cKO tendon vs. bone comparisons indicating less genes in similar pathways were differentially expressed in cKO animals compared to Ctrl. (E) Heatmap of top DEGs, clustered using Ward’s linkage, with a fold-change cutoff of 5 when compared to Ctrl tendon.

## Discussion

This study reveals that the tendon enthesis maintains a hypoxic niche during early postnatal growth. This unique microenvironment that persists beyond birth and underlies key aspects of fibrocartilage development, cell survival, and ECM organization. Through temporal mapping using EF5 staining, we showed that hypoxia is restricted to the mouse enthesis postnatally, while adjacent tendon and cartilage shifts towards higher oxygen concentrations soon after birth. This focal hypoxia highlights oxygen availability as a potential determinant of cell fate and ECM formation at the tendon-bone interface. Our findings build upon established parallels between the enthesis and the skeletal growth plate, both of which rely on hypoxia and HIF-1α for proper development.^3,11,13,14,24^ However, unlike the growth plate which fuses upon skeletal maturity, the enthesis maintains an “arrested” developmental state.^10,25^ In this study, we showed that loss of *Hif1a* in tendon and enthesis progenitors compromised enthesis cell survival during the postnatal growth, leading to impaired structural and cellular maintenance at the enthesis. This resulted in abnormal tendon-bone structural attachments, disrupted bone shape, and ultimately failure to maintain the fibrocartilaginous transition zone. These data suggest that maintaining HIF-1α is essential for the maintenance of a functional enthesis and is dispensable for tendon development and function.

The structural deficiencies we found in the enthesis and tendon are linked to underlying changes in TF biology. TFs are the primary architects of the collagen-rich tendon ECM, which is crucial for mechanical strength.^26^ During fetal development, tendon resident cells are highly proliferative, then shift to robust ECM production during postnatal growth.^27^ In other mesenchymal cells, hypoxia can induce chondrocyte differentiation, and positively regulate collagen deposition, yielding changes in ECM density that can further oxygen depletion.^13,28–30^ HIF-1α is a master regulator of proliferation and differentiation of chondrocytes,^13,14,31^ and prior studies have also shown that hypoxia can increase cell proliferation and limit differentiation with minimal changes in ECM production in human tendon-derived stem cells and neonatal porcine tenocytes.^32^ HIF-1α also regulates the deposition of osteochondrogenic matrix *in vitro*.^33^ We found that loss of *Hif1a* resulted in upregulation of genes associated with fibroblast differentiation, such as *Pi16*, *Dpp4*, *Fap*, and *Bmp4*, as well as *Crabp2*, regardless of oxygen tension, suggesting that *Hif1a* may act independently of hypoxia in maintaining a fibroblast progenitor fate *in vitro*.^34,35^ The premature differentiation of progenitors into fibroblasts may drive the aberrant deposition of proteoglycan-rich ECM in the mature cKO enthesis.

We further characterized the metabolic shifts typically associated with hypoxia. In other mesenchymal populations, hypoxia induces a switch from oxidative phosphorylation and mitochondrial respiration to glycolysis, a pathway essential for cell survival under low oxygen conditions.^36^ Chondrocyte survival in growth plate hypoxia, for instance, relies on the suppression of mitochondrial respiration.^17,37,38^ *In vitro,* Hypoxia can induce chondrocyte differentiation from mesenchymal progenitors and positively regulate collagen deposition, yielding changes in ECM density that can further oxygen depletion.^13,28–30^ Our transcriptomic profiling of TFs revealed that *Hif1a*-deficient cells were less capable of executing these metabolic switches, supporting the idea that hypoxic adaptation in the tendon requires HIF-1α. Interestingly, despite classical concepts, we did not observe increased TF proliferation or enhanced ECM deposition *in vitro* under hypoxic conditions, suggesting that tendon fibroblast responses to hypoxia may be more nuanced, and perhaps dependent on additional factors.

Further, our spatial transcriptomic data unveiled a possible dysregulation of cell proliferation and growth pathways at the enthesis in which the cells may be signaled to proliferate but are failing during the cell cycle due to DNA damage, possibly leading to the dysregulation of cell population homeostasis.

Pathologically, the disrupted cellular environment at the cKO enthesis may promote recruitment of macrophages and other immune cells, compounding tissue damage and degeneration.

Elevated HIF-1α in tendinopathy is associated with proinflammatory cytokine production and apoptosis, contributing to ECM breakdown and poor healing outcomes.^39^ Other oxygen tension-dependent mechanisms such as the modulation of *Rac1* activity have also been implicated in tendon disorders, reinforcing the clinical relevance of these developmental mechanisms.^28^ Further, the developing tendon and enthesis may experience a different oxygen environment than mature tendon^40^, and it is important to note that while HIF-1α can be induced by hypoxia, HIF-1α expression can also be induced by inhibiting the prolyl-4-hydroxylase domain (PHD) enzymes, which leads to the stabilization of HIF-1α.^41^ For example, transforming growth factor-β1 (TGF-β1) can inhibit PHD expression and stabilize HIF-1α in fibrosarcoma cell lines and mouse embryonic fibroblasts.^42,43^ As such, we found oxygen-independent mechanisms by which HIF-1α may also be regulated in the tendon and enthesis.

The observed modifications in gene expression patterns of tendon fibroblasts *in vitro* further support the hypothesis that HIF-1α modulates metabolic switches, ECM synthesis, and cytoskeletal organization under low oxygen tension. Specifically, the blunted transcriptional and metabolic responses of *Hif1a*-deficient TFs to hypoxia, along with changes in ECM-related gene expression, suggest that adaptation to hypoxia is not solely compensatory but may be required for proper ECM assembly and maintenance. We showed that the ability of tendon progenitors to deposit ECM was also dependent on *Hif1a*, and we have identified potential regulators of cell function including cell cycle and ECM regulation that were independent of hypoxia. The upregulation of apoptotic markers and alterations in cell cycle-related genes hint at possible mechanisms for impaired tissue integrity in the absence of *Hif1a*.

Taken together, our results indicate hypoxia as a defining feature of the developing enthesis, with HIF-1α functioning as a gatekeeper of cell survival, ECM organization, and tissue architecture at the tendon-bone interface. These discoveries have important implications for understanding enthesopathies, such as those seen in rotator cuff injuries, where impaired ECM organization and cell death are observed.^44^ Furthermore, these data suggest that maintaining a hypoxic niche or manipulating HIF-1α signaling could serve as a therapeutic or tissue engineering strategy to promote functional enthesis regeneration.

The maintenance of a hypoxic niche at the postnatal enthesis and the necessity of HIF-1α for cell survival and structural organization establish new principles underlying the development of the tendon-to-bone interface. Our results open avenues for further research and ultimately inform potential therapies for tendon and enthesis repair through the manipulation of oxygen in the interfacial environment to further influence the guided regeneration of the cellular and extracellular matrix gradients necessary for mechanical function post-injury.

## Materials and Methods

### Mouse husbandry

*Hif1a^flox/flox^; ScxCre* mice were generated by crossing male *Hif1a* heterozygous (*Hif1a^flox/WT^) ScxCre* mice with homozygous *Hif1a^flox/flox^* females to generate constitutive knockout mice (cKO; *Hif1a^flox/flox^*;*ScxCre*) and control mice (Ctrl; Cre-negative *Hif1a^flox/flox^* or *Hif1a^flox/WT^*;*ScxCre*). Mice were euthanized following IACUC approvals (carbon dioxide asphyxiation if >8 days of age or decapitation if <8 days of age). When possible, left and right limbs from each mouse were used in complementary assays (e.g., histology/nano-computed tomography using left limbs and biomechanical testing using right limbs) to reduce animal numbers. A total of n=156 mice, including controls, were used across all time points.

### EF5 staining

Due to the intrinsic link between hypoxia and *Hif1a*, we first determined if a hypoxic environment exists in the developing Achilles enthesis using EF5 [2-(2-nitro-1*H*-imidazol-1-yl)-*N*-(2,2,3,3,3-pentafluoro-propyl) acetamide] staining, which in the presence of hypoxia forms cellular adducts that can be visualized via immunostaining (n ≥ 3 C57BL6J embryos or mice per time point).^45^ 10mM 2-nitroimidazole EF5 compound was administered intraperitonially in timed-pregnant C57/BL6J dams (E16.5) or subcutaneously in neonatal pups 2hr prior to euthanasia (at postnatal (P) day 1 and P5). Hindlimbs were embedded in optimal cutting temperature (OCT) glycol-based embedding media, cryosectioned, and stained with ELK3-51 AF488 antibody (75μg/mL) with and without competed stains (Sigma, EF5-30A4M). Cells experiencing hypoxic stress accumulated EF5 adducts and were visualized using fluorescence microscopy (ECLIPSE Ni-U, Nikon, 20x). Tissues were counterstained with DAPI for nuclear staining.

### Gross anatomy, X-ray imaging, and forelimb grip strength

Ctrl and cKO mice were weighed at 4 or 8wks of age (n=15 for Ctrl and n=11 for cKO at 4wks of age; n=13 for Ctrl and n=8 for cKO at 8wks of age) and grip force was measured using a grip strength test device (BIO-GS4, Bioseb, Pinellas Park, FL). Mice were then sacrificed to visualize gross musculoskeletal anatomy and skeletal morphology using planar X-ray (Faxitron MX-20 (25kV, 8s; digitized with Fujifilm FCR Prima T2).

### Tendon and enthesis biomechanics

After euthanasia, intact left limbs from Ctrl (Cre-negative, n= 15; Cre-positive, n=3 males) and cKO mice (n=15) at 8wks of age were frozen at -20°C. Achilles tendons and calcaneus were dissected and kept hydrated (with removal of the plantaris tendon and surrounding muscles). Tendon volume was imaged using optical coherence tomography with polarization (TEL221PSC1, OCT-LK4 lens, ThorLabs Inc., Newton, New Jersey). Intensity and polarization images were acquired in 3D (A-scan averaging = 4; spacing = 10µm in x and y, 2.5µm in z; sampling rate = 28kHz) and intensity and retardation images were exported (32-bit TIFF). Slice analysis was used to determine tendon cross-sectional area (CSA) and bifurcation distance (Dragonfly, Comet Technologies Canada Inc., Montreal, Canada). After imaging, the proximal Achilles tendon was clamped in a thin film grip (Imada, Northbrook, IL, USA) with the calcaneus secured in a custom 3D-printed fixture (FormLabs 3B, Somerville, MA, USA) prior to uniaxial tensile testing. The tendon and grip were placed in a room temperature phosphate buffered solution (PBS) bath and a pin was used to secure the tendon-grip unit. Uniaxial tensile tests were conducted with load measured using a multi-axis load cell (±70 N; Mach-1 VS500CST, Biomomentum, Laval, Quebec, Canada). Samples were preloaded to 0.2N and preconditioned for 10 cycles (±0.025mm at 0.1 Hz). Grip-to-grip gauge length (L_0_) was measured at the start of each test after preload and was consistent between tests (3.6 ± 0.5 mm). Load and torque data were collected while applying displacement to failure at 0.01mm/min to assess axial and off-axis loading. Mechanical properties (e.g., maximum load, maximum strain, linear stiffness; linear modulus and maximum stress calculated using load and CSA) of the Achilles tendon were calculated from force-displacement data using a piecewise linear segmentation by dynamic reprogramming recursion package (dpseg) in R (v4.2.2 or later, The R Project for Statistical Computing, Vienna, Austria). Data were presented as mean ± standard deviation and compared using a one-way ANOVA and corrected for multiple comparisons using Tukey’s multiple comparisons tests (Prism v9.0+; Graphpad, LaJolla, CA, USA).

### Nano-computed tomography (Nano-CT)

Right mouse hind limbs at 8wks of age and fixed at 4% paraformaldehyde. After fixation, limbs were suspended in a 3% agar solution and scanned using a Nantom M scanner (nanoCT; 8 µm voxel resolution; 80 kV, 400 µA, 0.381-mm aluminum filter, 500-ms timing with 3 averaged frames and 1 skipped frame after axis rotation; 2,000 projection images acquired at a source-to-axis distance of 48 mm; Waygate Technologies LP, Pasadena, Texas). Three-dimensional reconstructed nano-CT images were analyzed using Dragonfly software (Version 2021.3; Object Research Systems, Inc, Montreal Canada) to measure calcaneal bone morphometry. Data were presented as mean ± standard deviation and compared using a one-way ANOVA and corrected for multiple comparisons using Tukey’s multiple comparisons tests (Prism v9.0+; Graphpad, LaJolla, CA, USA). For shape analysis of the calcanei, nanoCT images were filtered in Dragonfly using Laplacian smoothing (60 iterations) and STL files were imported into ShapeWorks Studio.^46^ Shapes were groomed to fill gaps and aligned based on anatomical landmarks and optimization (128 particles; initial relative weighting = 0.25; relative weighting = 75; starting/ending regularization = 1000; 1000 iterations per split; optimization iterations = 2000; Procrustes rotation/translation; Procrustes interval = 10; and narrow band = 4).

### Histology and TUNEL staining

Hind limbs from Ctrl and cKO mice at P7, P14, and P56 (8wks of age) were decalcified in 14% ethylene diamine tetra-acetic acid (EDTA, pH 7.2) after fixation and/or Nano-CT then paraffin embedded for histology. Tissues were embedded in the sagittal plane to visualize the Achilles enthesis and tendon midsubstance and sectioned at 5-10 µm and serial slides were stained for Hematoxylin and Eosin, Toluidine Blue, Safranin O, silver nitrate, or Picrosirius Red. Slides were imaged in brightfield using a 20x objective on a Cytation 10 (Agilent, Santa Clara, CA, USA). Sections stained with Picrosirius Red were imaged using circular polarized light microscopy with a 10x objective on an epifluorescent microscope (Leica) and a polarization camera and ThorCam software (ThorLabs Inc.) for quantitative polarized light microscopy (qPLI). The degree of linear polarization (DoLP) and angle of polarization (AoP) were analyzed using the Math and SciPy Stats libraries in Python3. Data were compared using 2-way ANOVA between genotype and age (Prism GraphPad, v10). Paraffin sections from Ctrl and cKO mice at P7 and P14 were stained using the In Situ Cell Death Detection Kit (Roche 11 684 817 910) with a DAPI (cell nucleus). TUNEL stained slides were imaged using epifluorescence microscopy (ECLIPSE Ni-U, Nikon, or Cytation 10).

### *In vitro* tendon fibroblast culture for gene expression and cell assays

Ctrl (Cre-negative) and cKO mouse tail tendon fibroblasts (TFs) were isolated from 4-6 month old male and female mice (n=3) for *in vitro* studies using the collagen gels (PureCol-S, Millipore Sigma).^47^ TFs from Ctrl and cKO mice were seeded on collagen-I-coated-tissue-culture plastic at 2,500 cells/cm^2^ in DMEM:F12 containing 1% FBS and 1% Pen/Strep at either 20% O_2_ or 1% O_2_ conditions (Whitley H35 Hypoxystation, Don Whitley Scientific, West Yorkshire, United Kingdom). Total RNA was isolated after 1-week in culture (for quantitative polymerase chain reaction, qPCR) or 4-days in culture (for bulk RNA-sequencing; RNA-seq) using spin-columns (PureLink RNA mini kit, Thermo Fisher Scientific) with on column genomic DNA digestion (RNase-free DNase, QIAGEN). RNA quality and quantity were checked using a NanoDrop spectrophotometer (qPCR; Thermo Fisher Scientific) or Bioanalyzer (RNA-seq; Agilent 2100 Bioanalyzer, Santa Clara, CA, USA). RNA with a 260/280 ratio greater than 2.0 was reverse transcribed to cDNA (qPCR; SuperScript IV VILO Master Mix, Thermo Fisher Scientific) and 10 ng/μL of cDNA was used per reaction. Quantitative real-time reverse transcription polymerase chain reaction (qRT-PCR) was performed in triplicate for each gene (Table 2) using a CFX96 Real-Time PCR Detection System (Bio-Rad) with Power SYBR Green PCR Master Mix (Thermo Fisher Scientific) with *Rplp0* and *Polr2a* were used as reference genes. After normalization to reference gene Cq values, fold change data were compared using two-way ANOVAs and corrected for multiple comparisons with Tukey’s multiple comparisons tests (Prism v9.0+; Graphpad, LaJolla, CA, USA).

**Table 2.**
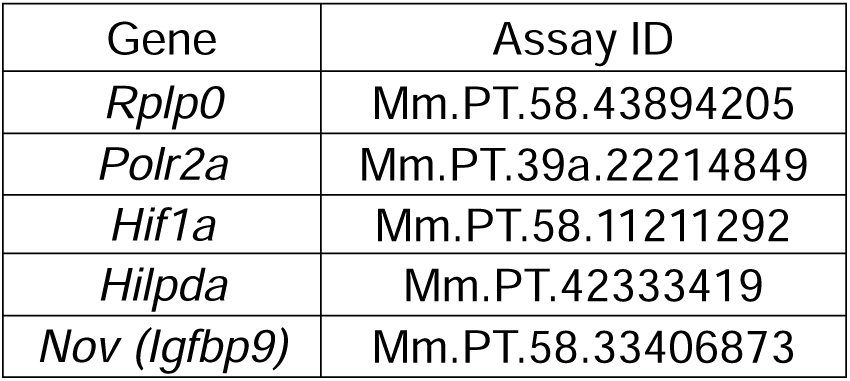
Integrated DNA Technologies (IDT) PrimeTime qPCR Primer Assay IDs.

| Gene | Assay ID |
| --- | --- |
| *Rplp0* | Mm.PT.58.43894205 |
| *Polr2a* | Mm.PT.39a.22214849 |
| *Hif1a* | Mm.PT.58.11211292 |
| *Hilpda* | Mm.PT.42333419 |
| *Nov (Igfbp9)* | Mm.PT.58.33406873 |

For RNA-seq (4-days in culture), samples (RNA Integrity Number (RIN) > 8.6) were submitted for Poly-A enrichment library preparation (Illumina) and sequenced using 151bp paired-end sequencing (Illumina NovaSeqXPlus, System Suite Version: 1.3.0.39308). De-multiplexed Fastq files were generated using BCL Convert Conversion Software v4.0 (Illumina). Cutadapt v4.8^48^ was used to trim reads and FastQC v0.11.8 was used to ensure data quality.^49^ Next, Fastq Screen v0.15.3 was used to screen for contamination.^50^ Reads were then mapped to the reference genome GRCm38 (ENSEMBL) using STAR v2.7.8a^51^ and assigned count estimates to genes with RSEM v1.3.3.^52^ Alignment options followed ENCODE standards for RNA-seq. Multiqc v1.20 compiled the results from several of these tools and provided a detailed and comprehensive quality control report.^53^ Differential gene expression was determined from count matrices with a paired design (e.g., WT v cKO) in DESeq2 in R/Bioconductor.^54^ We used the Database for Annotation, Visualization, and Integrated Discovery (DAVID) to analyze biological processes and Kyoto Encyclopedia of Genes and Genomes (KEGG) pathways.^55,56^

Additionally, mouse TFs from Ctrl and cKO mice were cultured for 2- or 4 days *in vitro* and ATP availability was measured in white opaque 96-well TC-treated plates (Falcon #353296) using CellTiter GLO 2.0 Cell Viability Assay (luminescence; Promega) per manufacturer recommendations. Data were analyzed using a two-way ANOVA with Sidak’s multiple comparisons for genotype and oxygen concentration (n=3 biological replicates with 1 male and 2 female for Ctrl and 2 male and 1 female for cKO; n=3 technical replicates per biological replicate). On parallel plates (96-well, black-walled, glass-bottom; CellVis #P96-1.5P), TFs were administered nascent metabolic protein labeling media^57^ with 1% FBS (replenished every 48hrs) to label nascent matrix deposited by TFs. After 4 days in culture, TFs were placed at 4°C for 5 minutes, washed with cold 1x Dulbecco’s Phosphate Buffered Solution (DPBS) then with cold 2% Bovine Serum Albumin (BSA) solution, and treated with 15μM DBCO-488 for 20-minute at 4°C. Following, TFs were washed with cold 2% BSA and fixed in 4% paraformaldehyde. TFs were stained with 0.1% CellMask Orange and Hoechst to label the cell membrane and nuclei, respectively. Lastly, TFs were imaged using spinning disc confocal microscopy at 20x strength on a Cytation 10 (Agilent, Santa Clara, CA, USA). Images were processed and analyzed using Gen5 (Agilent) to quantify nuclear size/shape and nascent matrix area. Data were compiled in GraphPad (Prism v9.0+; Graphpad, LaJolla, CA, USA). with two-way ANOVAs with Sidak’s multiple comparisons for genotype and oxygen concentration. Also in parallel, Ctrl and cKO cells were cultured in 20% O_2_ normoxia or 1% O_2_ hypoxia were cultured for two weeks and total GAG and calcium content from cell lysates were analyzed using a total glycosaminoglycan kit (abcam, ab289842), and a calcium assay kit (Sigma, MAK477), respectively.

### Spatial sequencing library preparation and analysis

All spatial sequencing data were generated using a recently published Seq-scope protocol.^58^ In brief, frozen glycol-embedded postnatal day 5 (P5) mouse hindlimbs were flash-frozen in liquid nitrogen and stored at -80°C until sectioning. 10 μm sections were mounted onto Seq-Scope Chip, processed from Illumina NovaSeq 6000 flow cell, and fixed in cold methanol. Tissues were stained with hematoxylin and eosin and imaged using a light microscope (Keyence, Itasca, IL, USA). Tissues were permeabilized with pepsin for 24 minutes to release mRNA onto the Chip. The library was generated and sequenced as described in the protocol.^58^ Data analysis was performed using a standard *novascope* pipeline as described in the protocol paper,^58^ to produce spatially indexed digital gene expression map (sDGE). Using sDGE and the polygonal coordinates drawn upon histology image matching sDGE, we segmented the insertional tendon and bone areas and extracted gene expression from the segments based on the spatial coordinates. Using the digital gene expression data, within-sample comparisons of tendon and bone (Ctrl and cKO samples) were performed as well as between-sample comparisons (Ctrl vs cKO tendon; Ctrl vs cKO bone). Differentially expressed genes were identified as genes with at least 5 reads in a sample with a p-value of 0.1 in comparisons within a section. Across sections, a fold change of 5 was used to identify differentially expressed genes. Counts were normalized to the number of reads within each ROI drawn. We used the Database for Annotation, Visualization, and Integrated Discovery (DAVID) to analyze biological processes and Kyoto Encyclopedia of Genes and Genomes (KEGG) pathways and R to analyze these data.

## Acknowledgements

*Hif1a* floxed mice were donated by Ernestina Schipani. We are grateful to Zachary Tata and Paige Cordts for conducting initial mouse husbandry and preliminary experiments, as well as LeeAnn Hold, PhD and Sophie Orr, PhD for consultation and experimental assistance. Histology and NanoCT were conducted with the generous support and talent of Emma Snyder-White, Carol Whitinger, and Andrea Clark in the Michigan Integrative Musculoskeletal Heath Core Center. Library prep and next-generation sequencing was carried out in the Advanced Genomics Core at the University of Michigan. Schematics were designed using Biorender.

## Disclaimers

The content is solely the responsibility of the authors and does not necessarily represent the official views of the National Institutes of Health, NSF, or NASA.

## Author Contributions

SSS, TP, and MLK contributed to conceptualization of the paper; SSS, TP, NM, ACA, and MLK contributed to design (methodology), SSS, NM, TP, SB, CL, SL, KL, ACA, and MLK contributed to investigation (performing experiments); SSS, NM, SB, SG, MS, AR, KF, CL, JHL, YS, ACA, and MLK contributed to analyses and interpretation (imaging and data analyses); SSS, NM, ACA, MK contributed to formal analysis of the findings being published; SSS and MLK contributed to drafting the article; All authors edited and approved submission of the article.

