## Supplemental material for "The developing tendon and enthesis are hypoxic and rely on hypoxia-inducible factor 1a (*Hif1a*) during postnatal development"

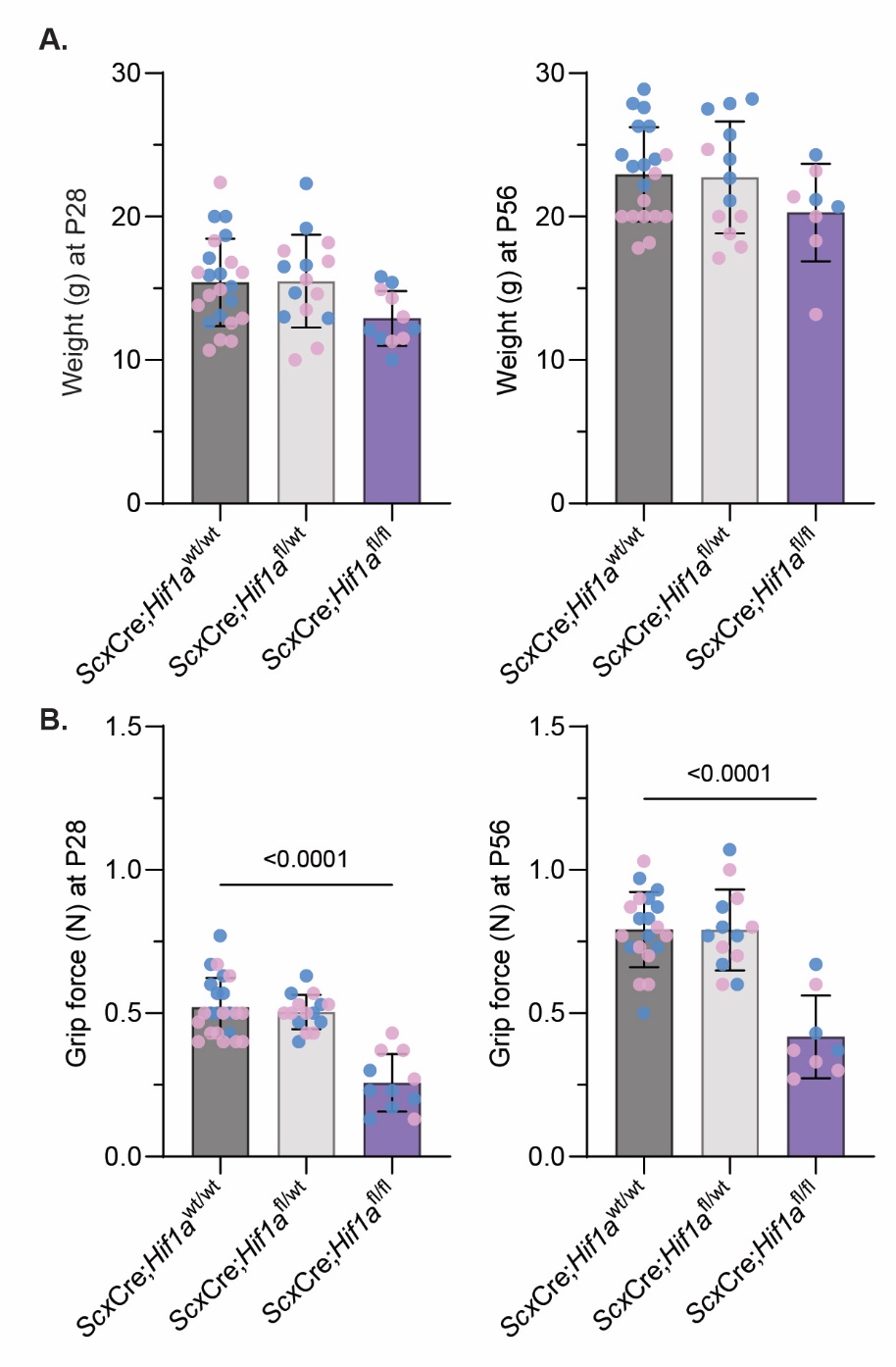

**Figure S1**. **Weight is comparable across genotypes while grip force is reduced in *Hif1a* knockout animals. (A)** Homozygous wild-type and heterozygous animals have comparable weights at P28 and P56. (B) Grip force is reduced in *Hif1a* knockout animals compared to homozygous and heterozygous controls and both P28 and P56 while homozygous wild-type and heterozygous animals have comparable grip forces at P28 and P56.

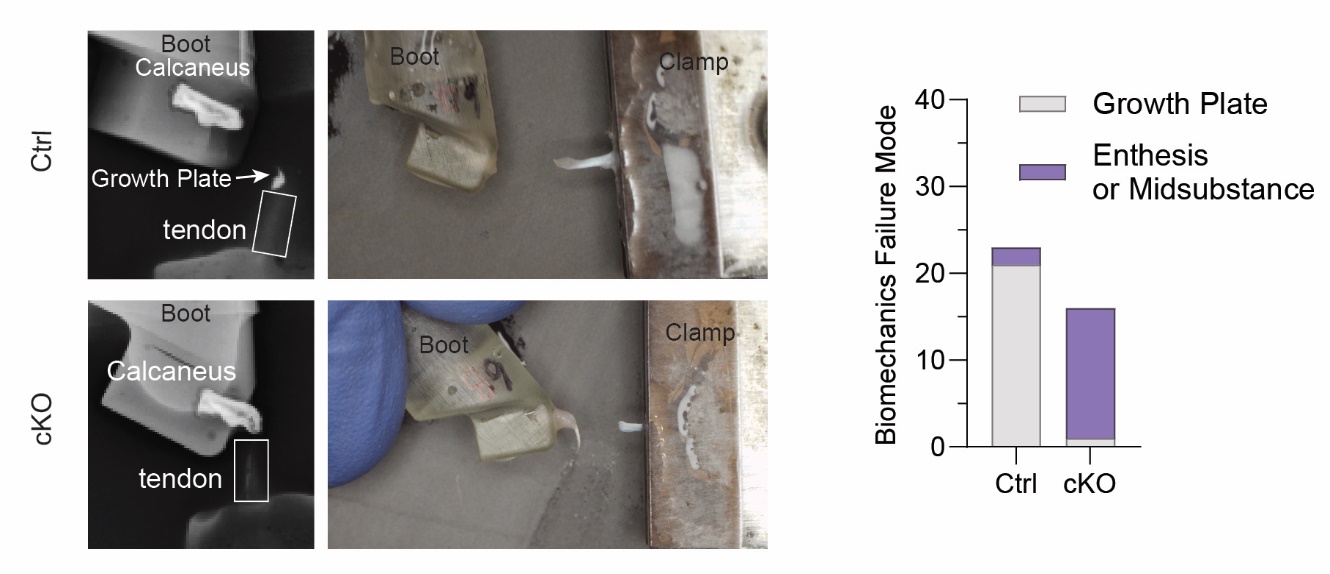

**Figure S2**. **Failure mode during mechanical testing of the Achilles tendon was genotype-dependent.** Representative X-ray images of Achilles tendons and calcanei in boots paired after biomechanics testing revealing failure mode in cKO tests were in the tendon and enthesis while Ctrl tests failed at the growth plate consistently.

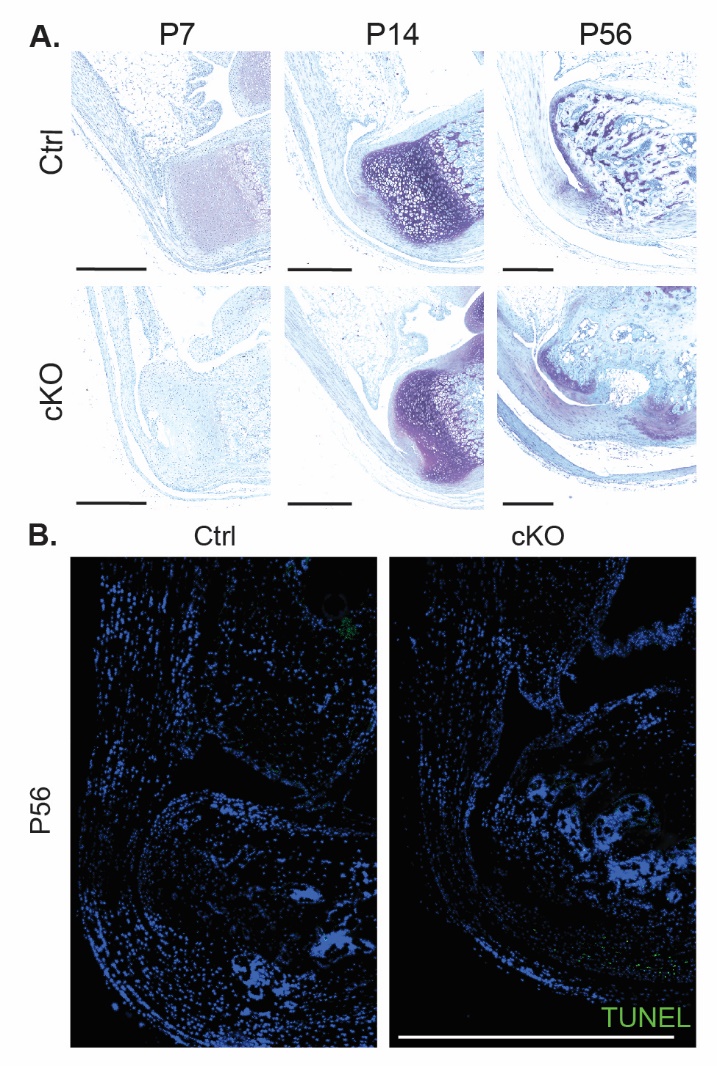

**Figure S3. Enthesis defects present during postnatal development.** (A) Hematoxylin and Eosin staining at P0 show unremarkable differences due to *Hif1a* knockout while disorganized cell and ECM are present at P7 and P56 entheses. (B) TUNEL staining at P56 revealed no apoptotic cells in the tendon and enthesis of Ctrl and cKO mice while some TUNEL positive cells were present at the base of the calcaneus in cKO mice. Scalebar = 1mm.

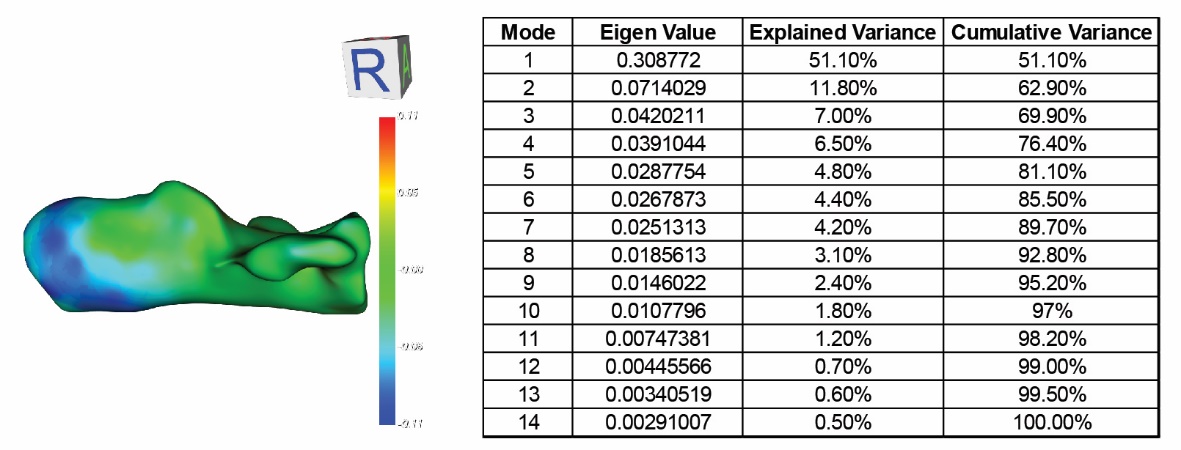

**Figure S4.** **Most of the differences in calcanei shape are due to changes near the tendon-to-bone interface.** Modes 1 and 2 account for over 60% of the change in shape.

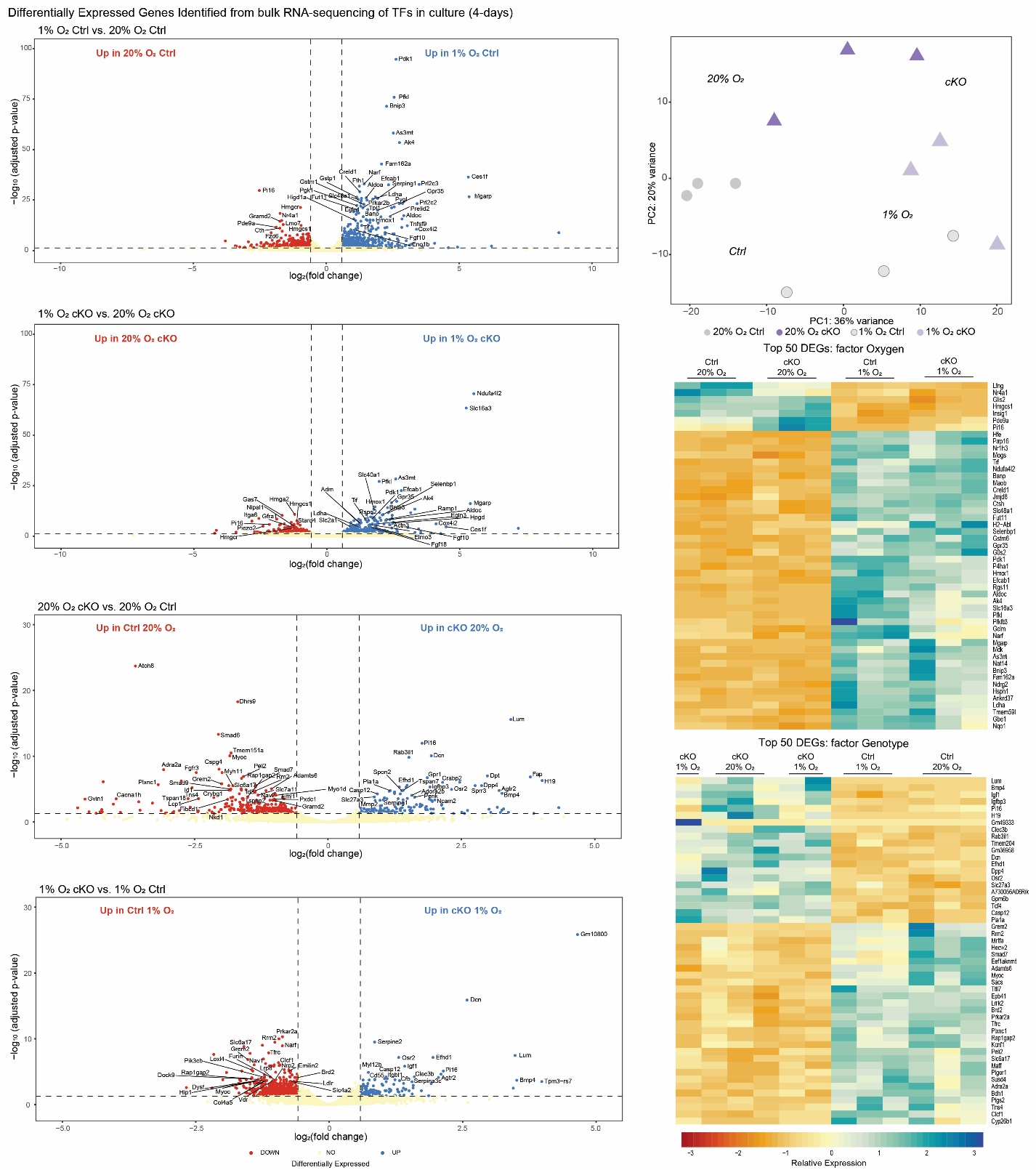

**Figure S5.** **Top 50 differentially expressed genes across all comparisons in 4-day RNA-sequencing data of *in vitro* cell cultures.**

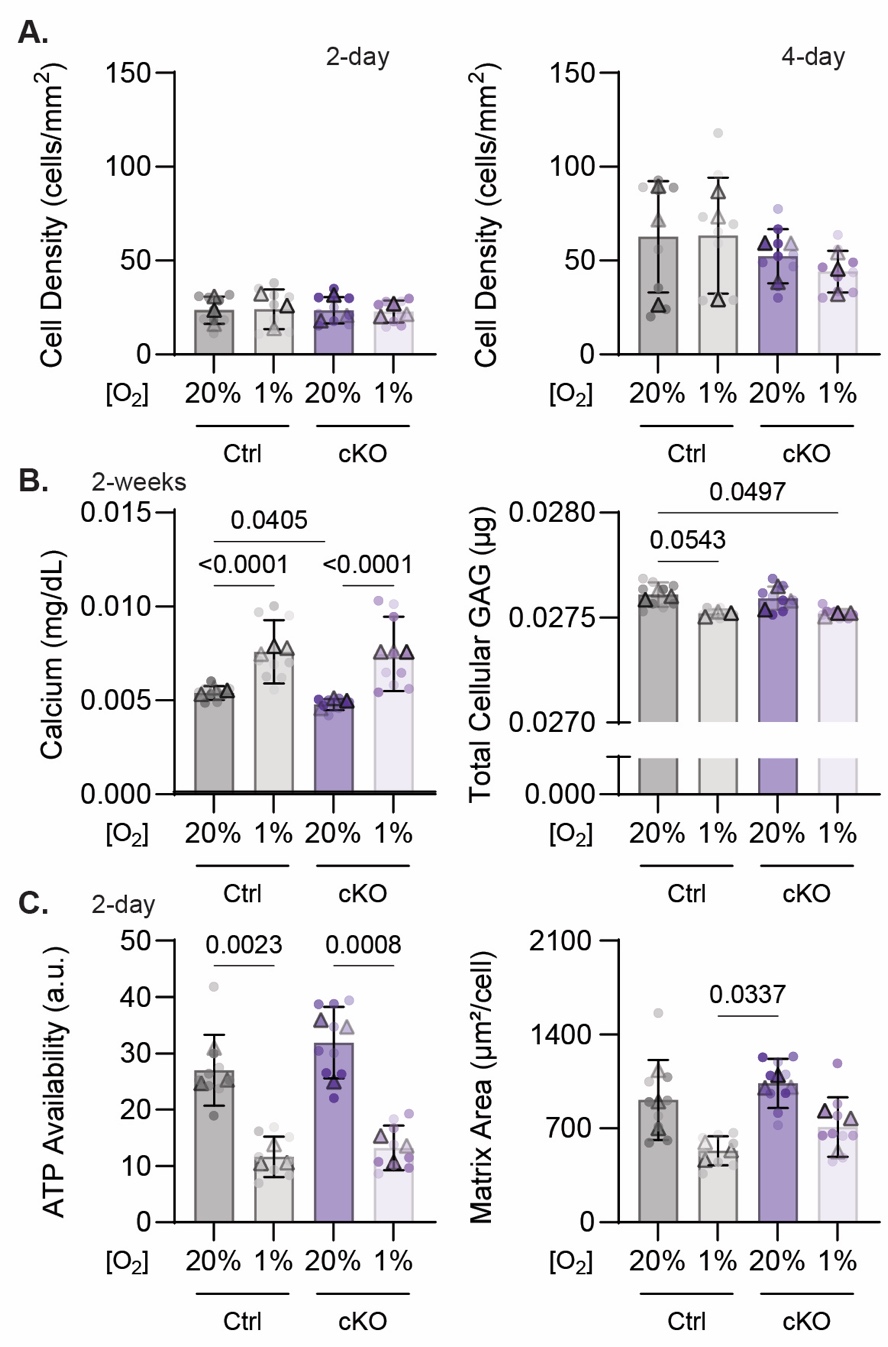

**Figure S6.** **Hypoxic conditions yield increased calcium and reduced GAG, ATP availability, and matrix area.** (A) Cell density did not change at either 2- or 4-days in cell culture across conditions while initial seeding density was similar. (B) Calcium available in solution is increased in hypoxic 4-day cultures while GAG from cell and matrix lysates is reduced at 2-weeks. (C) 2-day cultures show similar trends in ATP availability and matrix area as 4-day data with reduced ATP available and nascent matrix deposited in hypoxic conditions regardless of genotype (Figure 8). Triangles denote average of one biological replicate across 3 technical replicates.

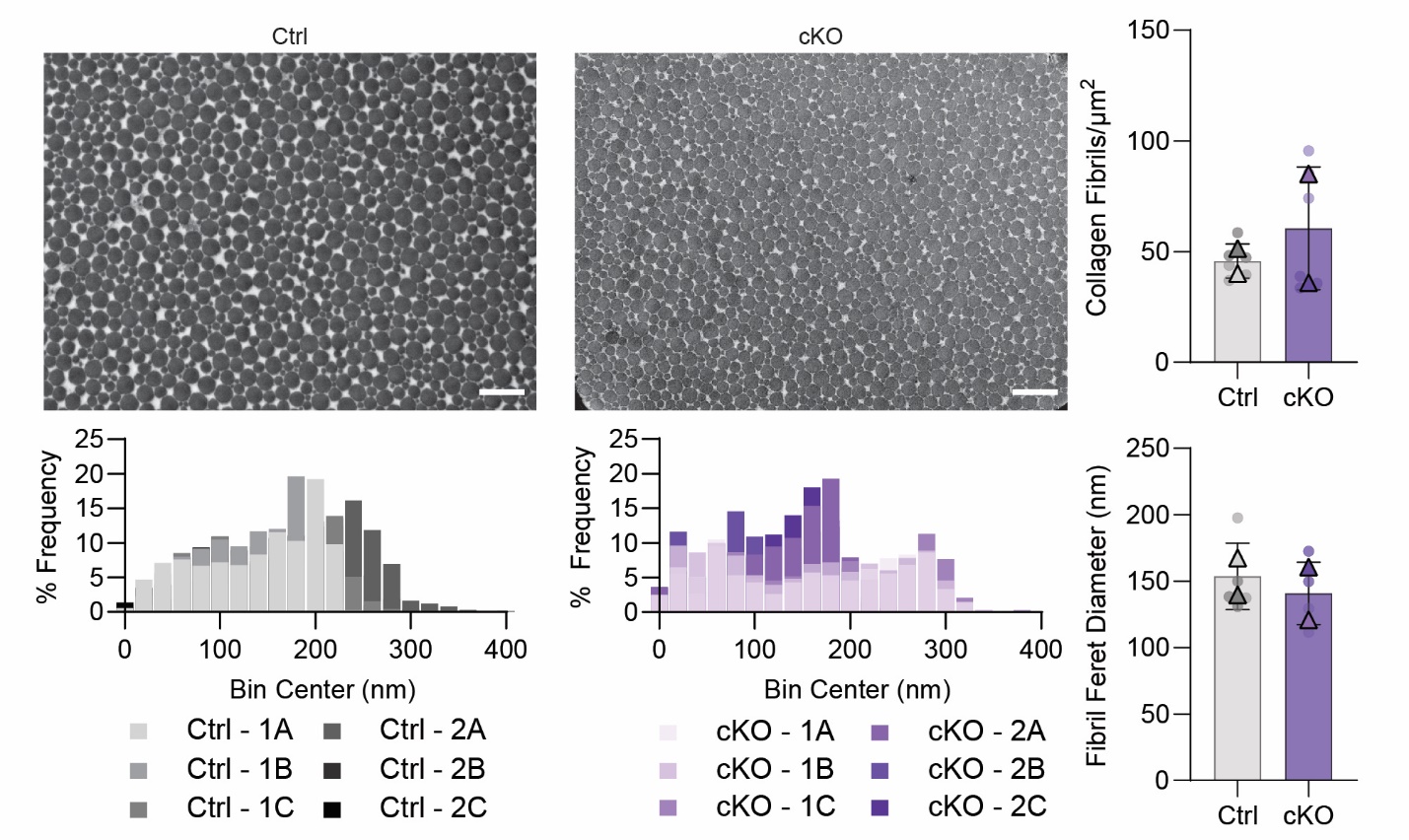

**Figure S7. TEM imaging of collagen fibrils in the Achilles tendons show a change in size distribution across genotypes.**

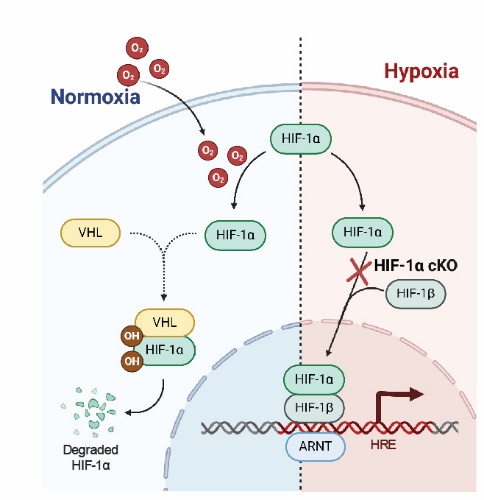

**Figure S8. Oxygen-dependent regulation of hypoxia-inducible factor 1α.** HIF-1α is recognized by the Von Hippel-Lindau (VHL) protein for hydroxylation and is naturally degraded under normoxic conditions (e.g., 20% O_2_). Under hypoxic conditions (e.g., 1% O_2_), HIF-1α is stabilized and translocated to the nucleus to enable transcription.
